# Shifting forage selection subsidizes seasonal resource scarcity

**DOI:** 10.64898/2026.03.13.711571

**Authors:** Jack G. Hendrix, Kristy M. Ferraro, Allegra E. Love, Jill M. Kusch, Dara Albrecht, Shawn J. Leroux, Quinn M. R. Webber, Eric Vander Wal

**Affiliations:** Cognitive and Behavioural Ecology Interdisciplinary Program, Memorial University of Newfoundland and Labrador, 45 Arctic Ave, St. John’s, NL, A1C 5S7, Canada; School for Environment and Sustainability, University of Michigan, 440 Church St, Ann Arbor, MI, 48109, USA; Department of Integrative Biology, University of Guelph, 50 Stone Rd E, Guelph, ON, N1G 2W1, Canada; Department of Biology, Memorial University of Newfoundland and Labrador, 45 Arctic Ave, St. John’s, NL, A1C 5S7, Canada; American University Washington College of Law, 4300 Nebraska Ave NW, Washington, DC 20016, USA

## Abstract

1. Nitrogen (N) is limiting for terrestrial herbivores, particularly over winter. Caribou (*Rangifer tarandus*) have adapted to seasonal scarcity of N by accruing muscle mass during the growing season when N is more abundant.
2. Nitrogen stored in muscle tissue is then relied upon during winter to compensate for dietary deficits. Once their diet shifts from N-rich vascular plants to N-poor lichen over winter, caribou can lose ∼30% of their muscle mass. As catabolized N is shed in urine on wintering grounds, caribou could act as elemental transport across seasons and landscapes. Furthermore, if deposited N is taken up by lichen or other winter forage, it might enrich the nitrogen-poor winter diet of caribou in the future.
3. We tested this potential transport via three steps. We analysed *Cladonia spp.* lichen and vascular plants upon which caribou forage across Fogo Island, Newfoundland, using %N content as our metric of forage quality. We then compared seasonal habitat selection responses to forage quality by caribou using integrated step selection analyses. In summer, caribou selected areas with higher vascular plant %N but did not select nor avoid *Cladonia* quality. In contrast, caribou selected sites with higher quality *Cladonia* in winter but responded neutrally to vascular plant quality.
4. We compared seasonal distributions of caribou to determine whether nitrogen consumed in summer and deposited in winter would occur in spatially discrete locations. Population-level kernel density estimates for summer and winter in this island herd were mostly non-overlapping, lending credence to the potential landscape effects of N transport.
5. When viewed together with established seasonal changes in woodland caribou physiology, sociality, and forage preferences, the shifts in habitat selection and seasonal ranges we observe here could serve as an adaptive strategy for caribou to recycle N and mitigate winter nutrient scarcity.

## Introduction

As stated in a 15th-century proverb, “*winter eats what summer grows*” (Meech, 1940). This adage captures a fundamental ecological constraint: organisms require consistent resources for bodily maintenance, survival, and reproduction, but resource availability is often inconsistent through time. Seasonal pulses of productivity therefore must be transformed into biological stores to sustain individuals through prolonged periods of scarcity. While humans and other food-caching species collect and store resources during periods of abundance (Andersson & Krebs, 1978), other wildlife must find alternative strategies to deal with periods of scarcity.

At high latitudes, short growing seasons and long winters create pulses of high-quality forage followed by long periods of resource scarcity (Cooper, 2014). Summer and early autumn are thus critical windows for animals to acquire forage and build body stores (Barboza et al., 2018), while winter is typically a period of conservation, during which minimizing losses of body material can be as important as acquisition itself (Barboza & Parker, 2006, 2008). Crucially, however, resource availability varies not only in time but also in space; animals must be in the right places at the right times to meet shifting energetic and nutritional demands. Therefore, characterizing the nutritional or elemental landscapes available to animals across time may help explain animal movement and, in turn, inform management and conservation. For northern species facing widespread declines (Cuyler et al., 2020; Smith et al., 2020), animal movement decisions that once aligned reliably with resource availability in space and time are increasingly decoupled by climate change (Post et al., 2009), habitat fragmentation, infrastructure, and other anthropogenic pressures. Without knowing when and where key nutritional resources are available, and how animals respond to these resources, it is difficult to identify critical seasonal habitats or movement corridors for animals navigating annual nutrient budgeting.

Nutritional and foraging ecology have historically considered how animals respond to variability in food resources in their environment. However, animals can also modify the nutritional environment themselves through behavioural and physiological processes (Hobbs et al., 1982). The field of zoogeochemistry incorporates animal effects into biogeochemical processes, treating animals as important mediators of elemental cycling and transport (Schmitz et al., 2018). Nutritional ecology and zoogeochemistry could be considered alternate perspectives on the same reciprocal interactions between animals and their environment. Animal-derived inputs can shape elemental distributions within ecosystems (McInturf et al., 2019), generating spatial heterogeneity in availability and structuring local biogeochemical patterns (Ferraro et al., 2022). Zoogeochemical processes like the decomposition of carcasses contribute to elemental landscapes and subsequently create local nutrient hotspots for foraging (Bump et al., 2009). Nutrient hotspots can be magnified when animal inputs are aggregated in time or space. Reproductive materials—such as natal fluids, egg masses, and neonatal mortality— can produce spatially concentrated and temporally synchronized elemental inputs on calving grounds or spawning sites (Ferraro et al., 2025; Naiman et al., 2002). Similarly, urine deposition during animal aggregation can concentrate nitrogen on the landscape, a key nutrient for plant growth (Ferraro et al., 2024). Across global grasslands, nitrogen enrichment through repeated ungulate urine inputs creates grazing lawns that reinforce feedbacks between movement, excretion, and forage quality (Coetsee et al., 2011; Day & Detling, 1990; Steinauer & Collins, 2001). Through consumption and excretion, animals both remove nutrients from their environment and add them back, but these processes need not operate at the same time and place.

Animal movement has long been recognized as a transport mechanism for elements to move across landscapes (Frank et al., 1994; Helfield & Naiman, 2001). While movement behaviour transports elements in space, internal physiology and metabolism can effectively transport elements in time. Bone growth and maintenance can sequester minerals for decades, both before and after death (Abraham et al., 2022; Coe, 1978). Anabolic tissue processes operate similarly by sequestering elements, but on shorter time scales; muscle catabolism during periods of food limitation may act as an important pathway of time-delayed nitrogen release to ecosystems (Holdo et al., 2007). In northern systems, winter resource scarcity causes animals such as ungulates to rely on stored body reserves (Barboza et al., 2018). Losses of muscle tissue result in elevated nitrogen excretion as catabolised nitrogen is shed in urine (Barthelemy et al., 2018; Boertje, 1985). Winter catabolism could increase nitrogen availability for plants growing on winter ranges, potentially creating patches of relatively high-quality forage at the time of year when forage nitrogen is most limiting (Parker et al., 2005). Seasonal migration or changes in sociality could further amplify the impact of nitrogen inputs, if nitrogen is transported from different locations and concentrated in aggregate on winter ranges (Lesage et al., 2000). By jointly assessing forage quality across areas of animal use and patterns of spatial behaviour, we can link where animals deposit elements to how those inputs shape forage quality across the landscape.

For herbivores inhabiting highly seasonal northern landscapes, where both forage availability and quality vary dramatically across space and time, successful reproduction and overwinter survival depend heavily on exploiting brief windows of high nutritional opportunity (Barboza et al., 2018; Barboza & Parker, 2006). Northern ungulates select high forage quantity and quality in the summer (Johnson et al., 2021) and build up body stores for winter. Winter forage tends to be particularly poor in nitrogen as green plants senensce (Chapin et al., 1980). Caribou (*Rangifer tarandus*) are unique amongst ungulates in adopting a lichen-dominated diet overwinter (Webber et al., 2022). While lichen is relatively high in carbohydrates, it does not contain sufficient nitrogen to meet nutritional demands (Boertje, 1990). While nitrogen content in lichen is generally low, even slight improvements in winter forage quality would nonetheless reduce reliance on body stores and leave animals in better body condition for parturition and lactation (Parker et al., 2005). Quantifying the variation in forage quality across the landscape, rather than relying on land cover as proxy for forage availability, would allow researchers to move beyond descriptive movement analyses to the mechanisms underlying caribou space use.

If woodland caribou preferentially select winter areas with higher forage nitrogen, what processes generate and maintain these nitrogen-rich patches? Woodland caribou exhibit many behavioural changes across seasons, with notable shifts in diet (Webber et al., 2022), habitat selection (Hornseth & Rempel, 2016), and sociality (Webber & Vander Wal, 2021). Body condition also deteriorates with winter diet, leading to body mass loss as fat or muscle (Tyler et al., 1999). Woodland caribou are typically dispersed during calving and then aggregate in autumn and winter (Bergerud & Page, 1987; Peignier et al., 2019). While site fidelity at the individual level is low in winter (Joly et al., 2021; Lafontaine et al., 2017), herds tend to occupy persistent, if broad, winter ranges (Brown et al., 2000; Ferguson & Elkie, 2004). Additonally, habitat selection patterns also tend towards more open areas in winter and more forested or sheltered areas in summer (Hendrix et al., 2025), such that even in non-migratory populations, the locations used by woodland caribou in summer and winter ranges tend to be distinct. The shift between disparate summer and winter locations parallels the physiological change from building muscle to catabolizing tissue. Woodland caribou therefore shift from a net intake of nitrogen in their summer range to a net deposition of nitrogen on winter grounds, transporting nitrogen through both space and time. Such transport could facilitate an adaptive strategy where winter muscle catabolism enriches winter forage and promotes seasonal site fidelity.

### Study objectives

To characterize the nutritional landscape available to caribou, we combined satellite and topographic data with forage sampling to create a stoichiometric distribution model (StDM, *sensu* Leroux et al 2017). We hypothesized that underlying geomorphology and biotic factors would lead to spatial variation in the elemental content of the primary plant and lichen species consumed by caribou. We predicted that forage species would differ in their responses to topography based on their ecology. For example, wetland-adapted species would have higher N content in low-lying sheltered areas, while alpine-adapted species would have higher N content in higher elevation and slope locations. Furthermore, we predicted that lichen nitrogen content would differ most from vascular plant species due to their unique nitrogen assimilation strategy that is not reliant on uptake from soil (Dahlman et al., 2004).

Our second objective was to examine how caribou movement and habitat selection varied across seasonal contexts. To do so, we combined our forage StDMs with data from GPS collared caribou to link seasonal resource dynamics and movement decisions. We modelled caribou behaviour via an integrated-step selection analysis (iSSA), testing whether habitat selection differed seasonally following changes in diet composition. We hypothesized that caribou select habitat based on access to higher nitrogen content in their forage. As their dominant forage changes across seasons, their habitat selection would also change based on what forage they were seeking. Specifically, during summer as vascular plants form the bulk of the diet, we predicted that caribou would select for areas with higher nitrogen concentration among vascular plant species but be less responsive to areas with higher lichen nitrogen concentration. In contrast, as lichen becomes a more dominant part of caribou diet overwinter (Webber et al., 2022), we predicted that caribou would select for areas with higher nitrogen concentration in lichen and cease responding to vascular plant nitrogen concentration.

Finally, to assess whether seasonal shifts in spatial behaviour could facilitate the transport and concentration of nitrogen from summer to winter ranges, we derived autocorrelated kernel density estimates (aKDE) from the GPS data for summer and winter. The relative size and overlap of these ranges would indicate whether caribou occupy distinct seasonal ranges and the degree of winter aggregation. We supplement this analysis by comparing the average nitrogen content of forage inside and outside the defined winter range, and by estimating the magnitude of catabolised nitrogen input based on population size and extent of winter aggregation.

## Methods

### Study area

Fogo Island is a small (∼238 km^2^) island off the northeast coast of Ktaqmkuk, or what is now known as Newfoundland, Canada. Woodland caribou were introduced to the island in the 1960s by the provincial government (Bergerud & Mercer, 1989) and now number approximately 150 individuals. Coyotes (*Canis latrans*) expanded their range onto Fogo within the last 20 years. Coyotes likely prey on calves in spring and summer (Huang et al., 2021), but their potential predation pressure on adult caribou remains in dispute.

### Spatial variation in nutrition (StDM)

We used a stoichiometric distribution modelling (sensu Leroux et al., 2017) approach to characterise how forage nutritional quality varied in space. Between June 11-14 2022, we sampled n = 104 locations across the homogenously granite-underlain northern half of Fogo Island (Brushett, 2014). Median distance between sites was 5.1 km (min = 110 m, max = 17.2 km; Appendix A, Figure A1). Our sampling period coincided with spring greenup, and all samples were collected within four days to avoid short-term phenological effects on nutrient content (testing of sampling date on %N and %C found no temporal differences from beginning to end of our sampling period). We also collected a small, but representative, set of plant and lichen samples in the winter that confirmed %N content of the evergreen plant species collected and *Cladonia* was relatively stable (Carstairs & Oechel, 1978; Chapin et al., 1980). Therefore, we applied the same %N values to winter movement data as well (see Appendix B for winter samples).

We generated random sampling points, stratified by availability of habitat type based on the province’s land cover classification system (Integrated Informatics Inc., 2014). At each location, we placed a 50 cm x 50 cm sampling plot and sampled from our eight focal forage species (Table 1) known to be consumed by caribou in Newfoundland (Bergerud, 1972). We collected four subsamples from separate individuals of each species present within the plot, but if only one individual was present, we took samples from different branches or stems of that plant.

**Table 1.** Caribou forage species in Newfoundland present on Fogo Island.

| Scientific name | Common name(s) | Sites sampled (% of 104) |
| --- | --- | --- |
| *Alnus incana* | Speckled alder | 22 (21%) |
| *Betula nana* | Dwarf birch | 14 (13%) |
| *Cladonia spp. (rangiferina, stellaris, uncialis)* | Reindeer lichen, star-tipped cup lichen, thorn lichen | 62 (60%) |
| *Empetrum nigrum* | Crowberry | 48 (46%) |
| *Kalmia angustifolia* | Sheep laurel, lambkill | 77 (74%) |
| *Trichophorum cespitosum* | Deergrass | 18 (18%) |
| *Vaccinium angustifolium* | Lowbush blueberry | 34 (33%) |
| *Vaccinium vitis-idaea* | Partridgeberry, lingonberry | 42 (40%) |

We focused on aboveground biomass in our sampling, analogous to what we expect caribou would consume. For *Cladonia* lichen, we collected entire thalli by the handful; for woody vegetation, we collected green leaves and did not include woody tissues in our samples for analysis (Thompson & Barboza, 2014). Samples were dried at 60 °C for >24 h to prevent nitrogen volatilization, homogenized using a Spex Sample Prep 5120 Mixer/Mill, and analysed for percent carbon and nitrogen using a Costech ECS 4010 Elemental Analyzer with a Conflo III interface at the Yale Analytic and Stable Isotope Center (New Haven, CT, USA). Nitrogen content, analogous to protein content in these analyses, is particularly limiting for caribou populations (Barboza & Parker, 2008), and its availability is often considered a key forage quality metric. As such, our analyses focus on %N as the measure of forage quality. Additional StDMs summaries for %C and for C:N ratio are included in Appendix A.

### Extrapolating nutrients to the landscape scale

After obtaining %N for our species samples at each location, we built a predictive model using multiple publicly available spatial predictors, detailed below. We prioritized within-sample predictive power over parsimony, as we wanted to explain as much variation and predict %N across Fogo Island as accurately as possible. As such we included every predictor we could that has been shown to influence plant stoichiometry in other Newfoundland research (Balluffi-Fry et al., 2020; Leroux et al., 2017). These included two land cover classifications: Newfoundland’s Sustainable Development and Strategic Science land cover system (Integrated Informatics Inc., 2014) and Canadian Forest Service land cover mapping (Hermosilla et al., 2022), both at 30m resolution. We also used a digital elevation model (DEM) at a 5 m resolution (Government of Newfoundland, 2020) and daily normalized difference vegetation index (NDVI) at a 250 m resolution (Natural Resources Canada, 2022). All rasters were scaled to the finest resolution of our predictor set, 5 x 5m courtesy of the DEM. That is, the same land cover class and NDVI value was applied to every 5 x 5m cell within the 30 x 30 m or 250 x 250 m pixel. From the DEM, we also calculated several topographic metrics, including slope (change in elevation across the focal cell; range 0 - 21°), aspect (cardinal direction slope faces, range 0 - 360°, where 0° = North), topographic position index (relative elevation of focal cell vs. the average elevation of the eight adjacent cells, range -0.71 – 0.88), and terrain ruggedness index (average difference in elevation between focal cell and each adjacent cell, range 0 – 5.68). While forage samples analysed here were from only one year, they were similar to data collected the subsequent year for a different study (Ferraro et al., 2024), and relative differences in forage quality in similar systems have been shown to be repeatable between years (Richmond et al., 2020). We thus developed an atemporal model and used this fixed nutrient landscape as predictor for multiple years of caribou movement data.

### Caribou data and iSSA

Between 2016 – 2023, the Newfoundland and Labrador Wildlife Division deployed GPS collars (Lotek Wireless Inc., Newmarket, ON, Canada) on adult female caribou on Fogo Island (2016-2019 deployments GPS4400M, 1250 g; 2022-2023 deployments Litetrack Iridium 420, 440g). Collars were redeployed on the same individuals wherever possible, prioritizing multiple years tracking the same animal over collaring as many individuals as possible. There was an average of 24 individuals collared per year during this period (min = 14, max = 30). Capture and collaring procedures were all conducted by Wildlife Division staff following government protocols.

GPS collars recorded positions every 2 hours. We calculated the straight-line distances between consecutive fixes (hereafter steps) as the basis for our integrated step-selection model (iSSA, Avgar et al., 2016) to test whether caribou responded to variation in forage quality across Fogo Island. We cleaned the GPS data to remove erroneous fixes and infeasible steps, including those in the ocean and displacements >30km between fixes. We further omitted steps from the southern non-granitic portion of Fogo Island (n = 79 804; 16%) over which we did not project forage stoichiometric values as per our sampling regime. After cleaning, there were n = 340 655 steps for n = 46 individuals for the entire period.

To test our specific hypotheses regarding seasonal changes in space use and forage quality, we separated our GPS data into biological seasons using climate and vegetation data informed by caribou behaviour and life history. We defined winter as 1 January – 1 March, the period with continuous snow cover > 75% across years (Love et al., 2025). To define our spring-summer season, we used VIIRS/NPP Land Cover Dynamics (Earth Science Data Systems, 2025) that analyses changes in NDVI through time and calculates inflection points in plant growth. This provides a date of peak spring greenup for each year from 2013 – 2022. The average date for Fogo Island over this period was 16 June, which also aligns with the end of parturition for most individuals and the beginning of lactation. We used an end date of 16 August, so that our summer and winter periods were the same length, and to avoid any inclusion of pre-rut autumn behaviour. These seasons are conservative, particularly for winter which likely extends both earlier and later, but we erred on the conservative side with our seasonal delineation as we were not data limited within these seasons. We ran the same iSSA model structure separately for winter and summer, with our final winter dataset including n = 54 816 steps among n = 41 individuals, and summer including n = 47 805 steps among n = 42 individuals.

Steps were characterized by two main parameters: step length, or the distance between subsequent points, and turning angle relative to the prior step (where 0 = linear movement and 180 = complete reversal of direction). We used the distribution of these parameters for each individual to generate random available steps, against which we could compare observed steps to test for selection or avoidance of features. We used n = 10 available steps for each observed step and included a random ID for this “cluster” in the model such that all statistical comparisons were within these clusters, to better account for a realistic ecological range of choices available to the individual at the step level.

### Environmental covariates in the iSSA

We used the stoichiometric distribution models we developed as predictive surfaces in our caribou habitat selection analyses. We included *Cladonia* %N on its own as predictor, and calculated the average %N from the seven vascular species’ StDMs as a metric of community-level vascular plant forage quality. Our StDM predicted surfaces were calculated at the finest scale of our set of predictors, which was 5m x 5m from the digital elevation model. We aggregated %N values to a 30m pixel, to match the land cover cells used to predict habitat selection in this population (Webber et al., 2024). Some locations, such as those within bodies of water and on roads, had missing values from the StDM as we did not predict %N into these land cover types. When summarizing %N, we treated these missing values as zeros to reflect the fact that they do not offer any forage opportunities.

Land cover classes were a key predictor of forage quality in our stoichiometric models, so we did not include categorical land cover again in our iSSA due to issues of collinearity. However, given extensive evidence in this population that habitat openness and forest cover are important predictors of caribou space use (Atkinson et al., 2025), we dichotomized land cover from the CFS system (Hermosilla et al., 2022) into forest vs. non-forest and calculated the average proportion of forest cover within a 100×100m cell for each point.

Our final iSSA models thus included %N in *Cladonia*, %N in vascular plants, and proportion forest cover as environmental covariates, with a binomial 0/1 response of randomly generated vs. observed steps, which we fit as a mixed Poisson regression following the Muff method (Muff et al., 2020). In addition to these covariates, we also included log(step length) and cos(turning angle), representing the movement component of the iSSA. We further included interactions between log(step length) and our three environmental covariates, to test whether selection or avoidance of certain parameters was a function of movement rate. If a habitat is beneficial because it is easier to travel quickly through, then selection for that habitat could increase along with step length, whereas selection at shorter steps could reflect higher quality foraging or safety habitat (Dickie et al., 2020). Finally, to account for individual variation in behaviour, we included random slopes for each predictor and interaction with individual ID. We evaluated goodness of fit of our iSSA models using a lineup protocol as per recommendations in the literature (Fieberg et al., 2024); within each season, we simulated 120-step segments using our model output, and plotted 19 of these segments visually against one randomly-selected 120-step segment from one of our individuals (Appendix C). When observers were unable to identify the real data amongst the simulations within this set of 20 trajectories, we concluded that our models successfully replicated realistic animal movements.

Parameter estimates from an iSSA output are difficult to interpret directly in an ecological context, as selection is an inherently context-specific phenomenon relative to availability, so coefficients in isolation are not as meaningful. We calculated relative selection strengths (RSS) for each of our predictors (Avgar et al., 2017) to visualize our model output in context. RSS is a ratio reflecting the probability of an animal selecting a specific habitat value over some constant reference value. This ratio is calculated for a range of biologically plausible values to see how selection changes. All other predictors in the model are held constant at their average or modal value. This results in a curve for a given predictor, the slope and shape of which indicate selection or avoidance. RSS is commonly log-transformed, so that selection for a habitat value is represented by a positive RSS and avoidance by negative RSS. For our reference levels, we used the average environment experienced by each animal, i.e., for a given individual, we calculated the mean(proportion forested) among their observed steps, and used this as the reference level in our calculations of RSS for forest cover. For each of *Cladonia* %N, vascular %N, and forest cover, we calculated RSS for each individual separately, as well as a population-level RSS including 95% confidence intervals from the iSSA output, as a measure of uncertainty around each response. Finally, based on the step length interactions with our three environmental covariates, we calculated RSS using the 10th percentile of step length, the median step length, and the 90th percentile of step length for each individual. Plotting these three RSS curves against one another shows how selection or avoidance for that habitat value changes with movement rate.

All analyses were conducted in R v4.3.2 (R Core Team, 2023) in a *targets* workflow (Landau, 2021), relying on packages including *amt* (Signer et al., 2019), *betareg* (Kosmidis & Zeileis, 2025), *data.table* (Dowle & Srinivasan, 2023), *ggplot2* (Wickham, 2011), *glmmTMB* (Brooks et al., 2017), *lme4* (Bates et al., 2014), *sf* (Pebesma, 2018), *terra* (Hijmans, 2025), and *tidyverse* (Wickham et al., 2019).

### Potential nitrogen transfer from summer to winter ranges via catabolism

Despite being non-migratory, based on our field observations Fogo caribou appeared to use disparate locations in summer and winter. To validate this assumption, we used GPS collar data within the same seasonal date thresholds as described above in the iSSA and created seasonal utilization distributions for the population. We pooled all individuals and calculated autocorrelated kernel density estimates (aKDEs) (Fleming et al., 2015) for summer and winter and converted these to raster surfaces with 25 x 25m cells. We then calculated the correlation between the two seasons at each cell. A negative correlation between summer and winter aKDEs would indicate that high-intensity of use summer areas were not in the same location as high-intensity of use winter areas.

We also compared the %N content at our sampling locations based on whether they fell inside or outside the 50% winter kernel. If forage within this core area was relatively high in N, it could indicate that the presence of caribou enriches forage. Conversely it could also be an artifact of caribou selecting locations with more %N available. While causation cannot be determined from this analysis, it could support our assumption.

To more directly assess how caribou might influence their environment, we used previously published estimates of body mass and condition to calculate the amount of nitrogen associated with seasonal body mass loss in caribou. The amount of nitrogen excreted via catabolism, and thus the potential nitrogen subsidy moved from the summer to winter range, could be divided by the area of the winter range to get a nitrogen-per-area estimate. Field observations of the Fogo Island caribou population indicate an approximate 100:50 cow-to-bull ratio, consistent with other caribou herds (Brown et al., 2000). We collected data from previously published research on changes in muscle mass for different caribou populations from fall to spring (Adamczewski et al., 1987; Gerhart et al., 1996; Huot, 1989; Reimers et al., 1982) (Appendix E), particularly other island herds, to calculate average initial autumn muscle mass and the average percent loss of muscle through to spring. Estimates of catabolism among wild male caribou are lacking, but analyses of captive male reindeer indicated that males had approximately 35% higher muscle mass, as expected from allometric relationships of muscle mass and total body mass (Gerhart et al., 1996). We thus took the average change in muscle mass observed in female caribou and multiplied it by 1.35 to approximate male protein catabolism overwinter. We then estimated nitrogen loss by applying the standard conversion that elemental nitrogen comprises 16% of crude protein by mass, in line with previous work on elk (Hobbs et al., 1982) and wildebeest (Holdo et al., 2007, 2011), likely also applicable to caribou (Ferraro et al., 2022).

We used the 50% kernel of the winter aKDE as the area over which we estimated catabolized nitrogen deposition by caribou, to focus on the more densely occupied core portion of the winter range. Although the 50% kernel does not actually mean that 50% of the GPS points fall within this area, nonetheless caribou also spent time outside this area overwinter so that not all the catabolised nitrogen would be deposited within this core range. We thus approximated that 50% of the nitrogen deposition would occur within the 50% kernel. We multiplied the total nitrogen loss by 0.5 before dividing by the area of the core winter range to obtain an estimate of deposition in kg N/km^2^.

## Results

### Forage nitrogen content

Nitrogen content in caribou forage on Fogo Island varied both within and between species (Fig 1, Table 2). Deciduous shrubs (alder, blueberry) and graminoids tended to have higher %N, while crowberry and partridgeberry had relatively low %N, with *Cladonia* lichens having the lowest %N overall (Table 2). The explanatory power of our landscape model in predicting percent nitrogen varied by species (Table 2). Forage species differed in how topographic and land cover variables influenced their nutritional content (see Appendix A Table A1 for species-specific model summaries). For example, topographic slope was negatively related to %N in alder and birch samples, but a positive effect on *Cladonia*, sheep laurel, and lingonberry. Topographic position index (TPI) also had a negative effect on %N in alder, but a positive effect on sheep laurel, lingonberry, and birch samples. When the average vascular plant %N was calculated for each cell, there was a slight negative correlation with *Cladonia* %N (r = -0.058) in our StDMs. However, when considering pairwise correlations between all species, this pattern was more complex (Appendix A Table A3). While our StDM surfaces are imperfect descriptions of forage quality, they are comparable to previous %N StDM work in Newfoundland. Our pseudo-R^2^ for *Cladonia* nitrogen was 24%, and the average for vascular nitrogen was 40%, comparable with 31% (Balluffi-Fry et al., 2020) or 32% (Richmond et al., 2020).

**Figure 1.**
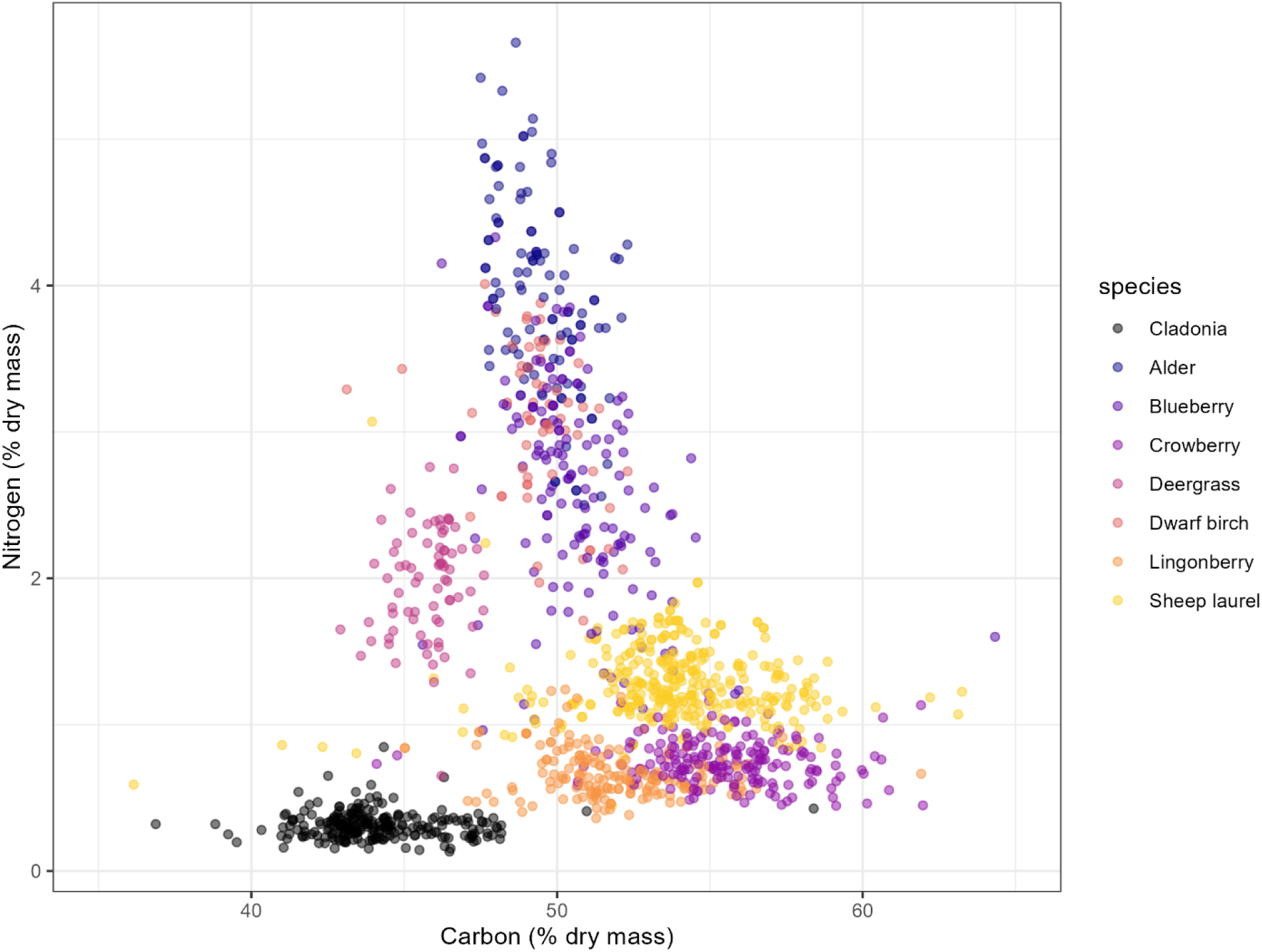
Percent carbon and nitrogen content of samples collected from eight caribou forage species on Fogo Island, NL. Of *n* = 1343 total samples, *n* = 11 outliers have been omitted from this plot for visualization purposes. Our analyses focused primarily on % nitrogen, but further information about % carbon and C:N ratios are provided in Appendix A.

**Table 2.** Average percent nitrogen content of forage species samples collected on Fogo Island, NL, June 11 –14 2022, along with model pseudo-R^2^ for predicting %N based on landscape covariates. The same model was applied to each species* and included Canadian Forest Service land cover, provincial land cover, NDVI, elevation (m), slope (%), aspect (°), distance to coast (m), topographic position index, and terrain ruggedness index. * CFS land cover and terrain ruggedness were removed from the dwarf birch models for convergence.

| Species | $n$ samples | % nitrogen ± SE | Pseudo-$R^2$ of N model (%) |
| --- | --- | --- | --- |
| Alder | 111 | $3.95 \pm 0.06$ | 63.6 |
| Blueberry | 147 | $2.61 \pm 0.06$ | 44.1 |
| *Cladonia* | 248 | $0.32 \pm 0.01$ | 23.9 |
| Crowberry | 190 | $0.76 \pm 0.01$ | 19.9 |
| Deergrass | 76 | $1.96 \pm 0.04$ | 36.4 |
| Dwarf birch | 60 | $3.04 \pm 0.07$ | 37.9 |
| Lingonberry | 177 | $0.67 \pm 0.01$ | 47.1 |
| Sheep laurel | 334 | $1.28 \pm 0.01$ | 31.6 |

### Caribou movement and habitat selection

Our iSSA models effectively predicted caribou movement. When presented with a lineup protocol for each of summer and winter, 1/9 participants successfully identified the real data amongst 19 simulations for summer, and 0/9 in winter (Appendix C). Model summaries are provided in Table 3.

**Table 3.** Parameter estimates from summer and winter integrated step-selection analyses of caribou on Fogo Island, Newfoundland. Summer model includes n = 519 105 steps in 47 805 clusters and 10 years over 42 individuals. Winter model includes n = 638 600 steps in 58 416 clusters and 9 years over 41 individuals. Model summaries are included for transparency, but note that for the purposes of interpretation, relative selection strengths (RSS) better reflect biologically relevant effects than directly interpreting beta coefficients (Avgar et al., 2017).

| | Estimate ± SE | $z$ | $p$ |
| --- | --- | --- | --- |
| Summer |  |  |  |
| log(step length) | $-0.121 \pm 0.010$ | -12.0 | < 0.001 |
| cos(turning angle) | -0.211 ± 0.007 | -30.73 | < 0.001 |
| Proportion forest cover (%) | 0.783 ± 0.087 | 9.06 | < 0.001 |
| *Cladonia* % nitrogen | -0.751 ± 0.150 | -4.99 | < 0.001 |
| Vascular % nitrogen | 0.093 ± 0.027 | 3.41 | 0.007 |
| log(step length) : forest cover | -0.028 ± 0.021 | -1.35 | 0.18 |
| log(step length) : *Cladonia* %N | 0.043 ± 0.036 | 1.221 | 0.22 |
| log(step length) : vascular %N | 0.043 ± 0.006 | 6.78 | < 0.001 |
| Winter |  |  |  |
| log(step length) | -0.020 ± 0.011 | -1.81 | 0.07 |
| cos(turning angle) | 0.016 ± 0.006 | 2.59 | 0.01 |
| Proportion forest cover (%) | 0.005 ± 0.201 | 0.025 | 0.98 |
| *Cladonia* % nitrogen | -0.822 ± 0.187 | -4.39 | < 0.001 |
| Vascular % nitrogen | 0.330 ± 0.029 | 11.56 | < 0.001 |
| log(step length) : forest cover | -0.436 ± 0.030 | -14.73 | < 0.001 |
| log(step length) : *Cladonia* %N | 0.441 ± 0.032 | 13.99 | < 0.001 |
| log(step length) : vascular %N | -0.043 ± 0.005 | -8.00 | < 0.001 |

Caribou on Fogo Island generally avoided more forested areas during winter and weakly selected more forest cover in summer, although there was substantial individual variation in these patterns (Appendix D). Forest avoidance in winter was stronger when caribou were moving more quickly (Table 3), as would be expected as more closed habitats tend to limit movement rate (Dickie et al., 2020).

### Caribou response to StDMs

The predicted vascular plant and *Cladonia* %N surfaces influenced caribou habitat selection patterns in summer and winter as predicted. While the model parameters appear to indicate a negative response, e.g., the coefficient for *Cladonia* %N in winter is negative, this is due to the interaction with step length term that is also included. When relative selection strength is calculated using the median step length, the selection for *Cladonia* %N overwinter is clearly positive (Fig 2). Indeed, when RSS is calculated for the 10^th^, 50^th^, and 90^th^ percentiles of step length, winter selection for *Cladonia* %N is only ambiguous at the very slowest movement rates of ∼4 m/h (Appendix D Fig D5).

**Figure 2.**
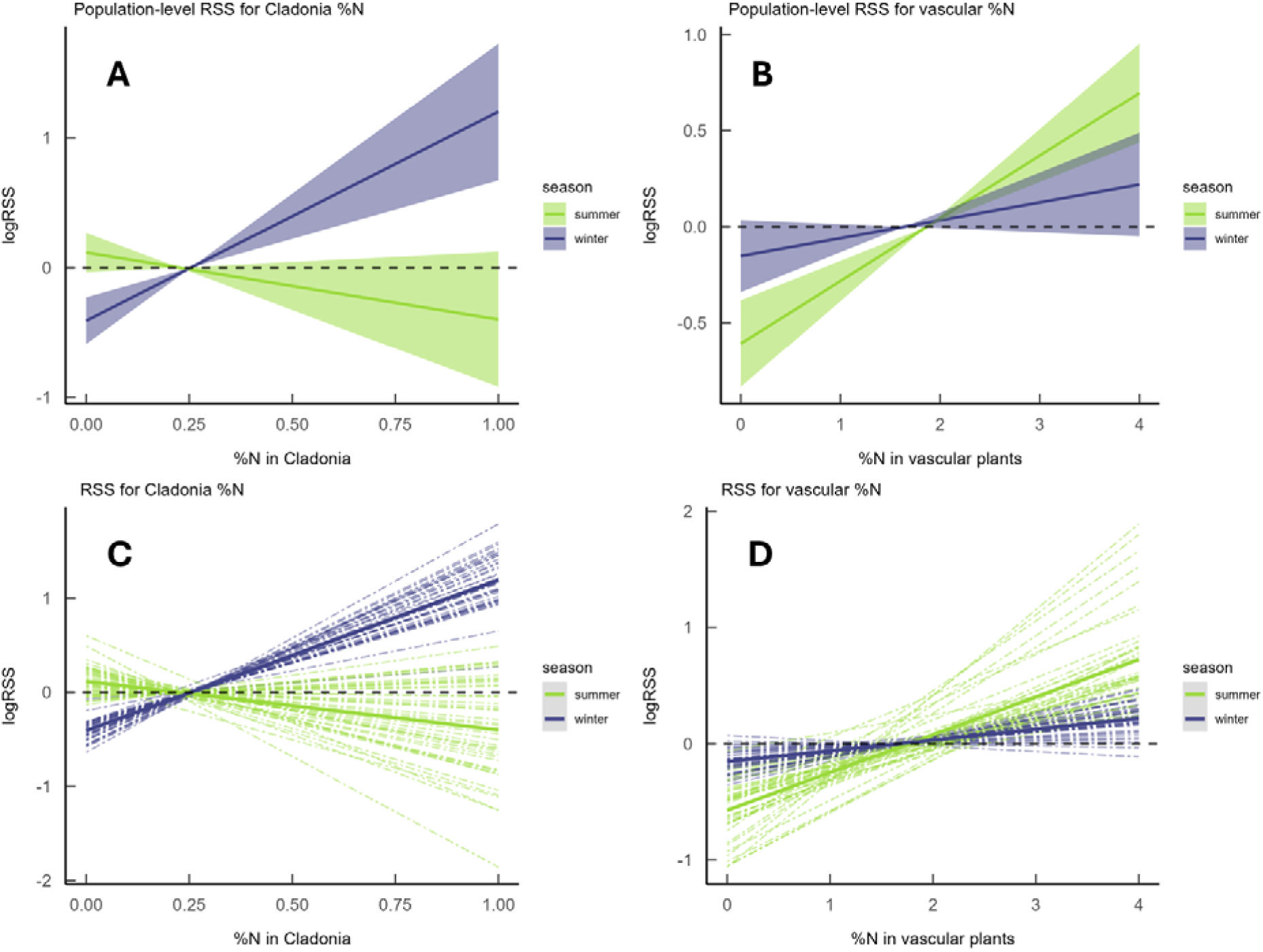
Relative selection strength (RSS) for vascular and *Cladonia* %N content, separated by season. (A) and (B) show population responses, with the shaded ribbon reflecting the 95% confidence intervals as calculated from the standard error of the parameter estimates for each predictor variable. Blue lines reflect winter (1 January – 1 March) movement and green reflects summer (16 June – 16 August). (C) and (D) show the same RSS, but with each individual given its own dotted line relative to the population estimate. logRSS values > 0 indicate selection for that x-axis value relative to the average %N, which is where the lines cross the x-axis. Note that the slope and intercept of individual lines differ more in summer than in winter, particularly so for vascular %N (panel D), reflecting the greater individual variation in selection behaviour for this season and predictor.

When interpreting relative selection strengths (RSS), values > 0 indicate that the study species is more likely to select that covariate value relative to the reference value. The reference in all RSS presented here were the average value of %N for that forage. Caribou selected for areas with higher %N content in vascular plants during the summer, when vascular plants are at their most nutritious, but were either neutral or weakly avoided areas with high *Cladonia* %N during the summer, when lichen is not a major diet component (Fig 2B). While there was some individual variation in these selection patterns, particularly with respect to vascular %N in summer (Fig 2D), these seasonal trends were robust to the 95% CI from parameter estimates (Fig 2A,B).

### Seasonal kernel density estimates and estimated nitrogen inputs via catabolism

Caribou on Fogo Island occupied distinct summer and winter ranges, particularly with respect to the highest intensity of use areas (Fig 3). Increased aggregation and sociality in winter resulted in one concentrated “hotspot” across the entire population, while dispersion during the summer resulted in several smaller high-intensity areas spaced out from one another. This aggregation vs. dispersion is reflected in the relative size of the 50% kernels: the summer 50% kernel land area was 64.1 km^2^ (95% CI: 44.5, 87.8) vs. 38.4 km^2^ (15.7, 76.0) in winter. The overlap between the 50% kernels for each season was 18.8%. Within the northern extent of Fogo Island where our StDMs and iSSA were conducted, continuous summer and winter kernel density were negatively correlated with r = -0.22.

**Figure 3.**
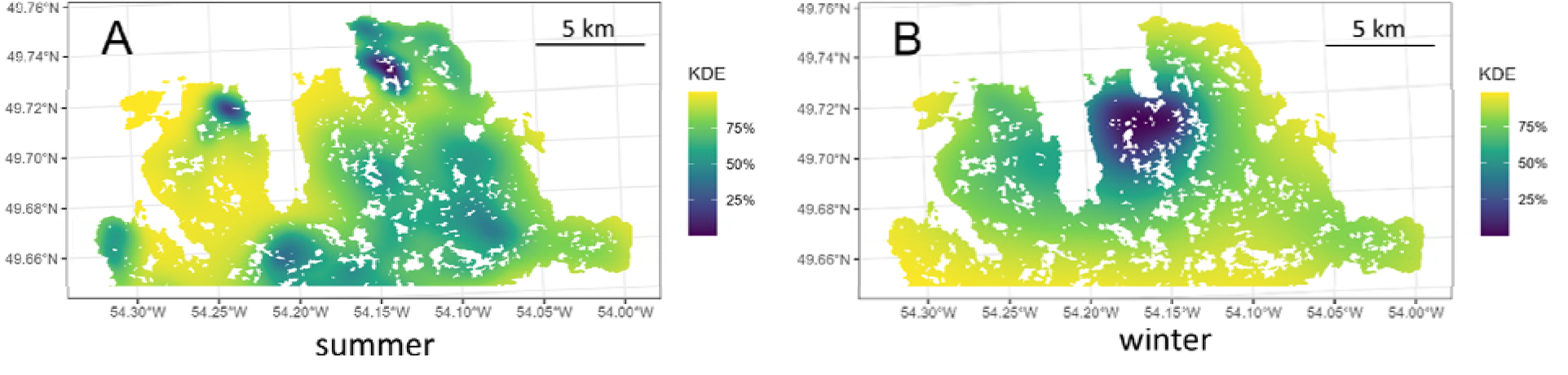
Population-level autocorrelated kernel density estimates (aKDE) for Fogo Island caribou in (A) summer (16 June - 16 August) and (B) winter (1 January - 1 March). The colour scale represents the % kernel, i.e. darker values represent more intensely used areas, with yellow areas on the edges of the utilization distribution.

Our comparison of forage %N at sampling sites revealed that crowberry and lingonberry samples inside the winter range of caribou had significantly higher %N than those outside, while alder showed the opposite effect with lower %N inside the winter 50% kernel. All other species showed a non-significant trend towards having slightly higher %N inside the winter range (Figure 4).

**Figure 4.**
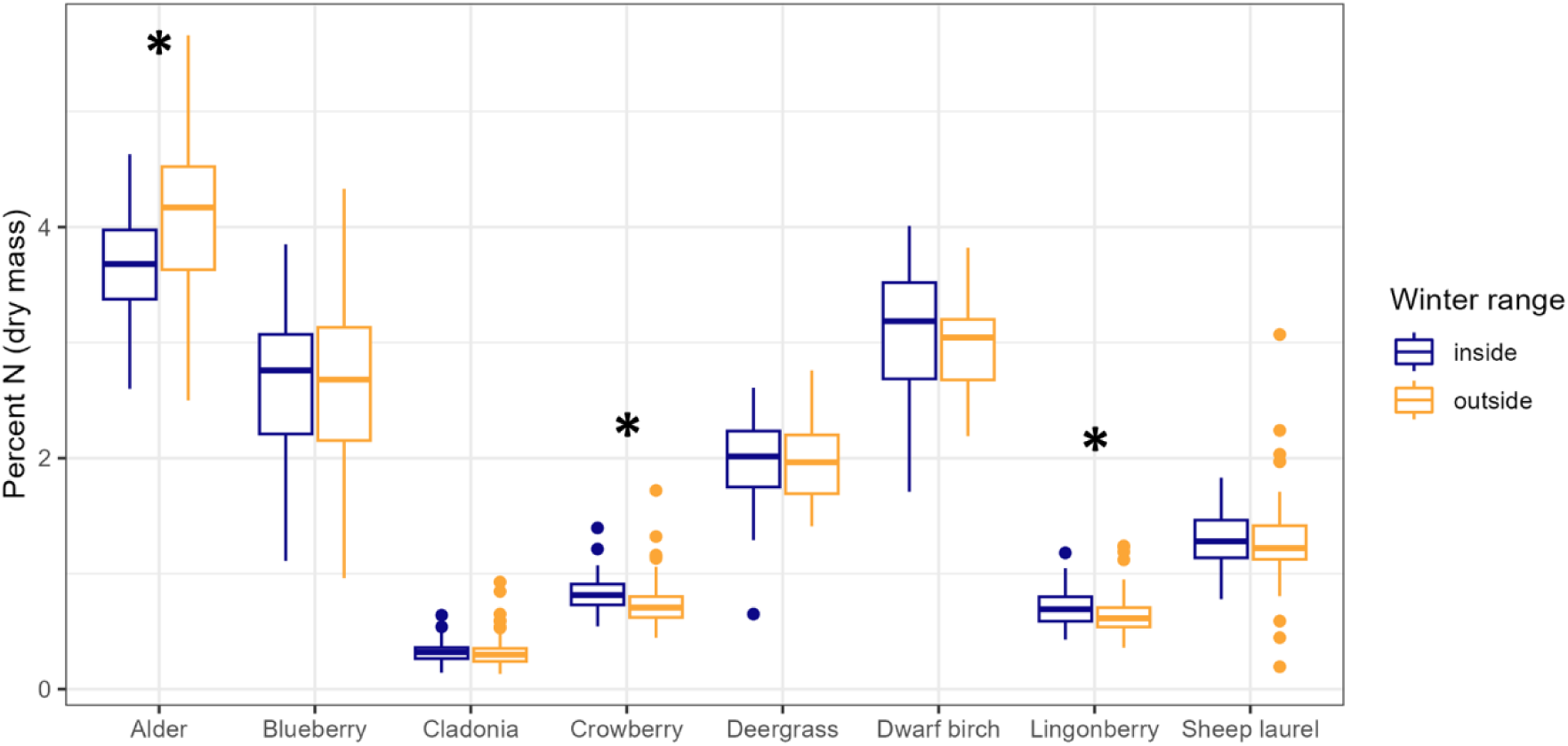
Comparing %N content of forage species sampled from within and without the 50% kernel of caribou overwinter on Fogo Island, NL. Asterisks **\*** indicate significant differences within species (p < 0.05).

Additionally, our estimates of catabolised N deposition suggest that a population of 150 caribou on Fogo Island may deposit 170 kg of N within their winter grounds via catabolism. With the 50% kernel containing 38.4 km^2^ of land, and assuming only 50% of catabolised N is deposited within this area, the population could be adding 2.2 kg N/km^2^ (see Appendix E for full calculations). Atmospheric deposition data are not available for Fogo specifically, but recent estimates for elsewhere in NL suggest atmospheric N deposition is approximately 0.2kg/ha, or 20kg/km^2^ (Cheng et al., 2022). The N deposited by the caribou population overwinter may thus be ∼11% of annual atmospheric deposition in the same area.

## Discussion

Here we combined fine-scale seasonal habitat selection patterns with a quantitative map of forage quality to assess the potential for adaptive nutrient transport by ungulates. Using an integrative approach, we show that caribou move through space in seasonally structured ways to meet nutritional demands, responding not only to the presence of forage but to spatial variation in forage quality. In summer, individuals select areas where vascular forage has higher nitrogen concentration, allowing them to capitalize on periods of high resource quality to build nitrogen stores. In winter, animals shift their space use toward areas where lichen nitrogen concentration is highest, reflecting both a transition to the dominant available resource and selection for relatively higher-quality forage. Such seasonal shifts likely represent a loss-minimizing strategy during periods of nitrogen scarcity. Together, these season-specific movement strategies reveal a spatial mechanism by which animals track high-quality resources through time in parallel with changes in diet.

Caribou diets are varied, and even while winter diets are predominantly lichen, they continue to consume other forage sources (Webber et al., 2022).We were nonetheless able to detect clear season-specific selection for %N in forage as predicted, indicating that resource selection analyses are robust to assumptions about which forage species animals consume and where those forages occur across the landscape. Similar results have been reported in Newfoundland and elsewhere, where simplified forage-quality models based on one or a few key species successfully predicted animal space use despite incomplete knowledge of diet composition and forage distributions (Balluffi-Fry et al., 2020; Richmond et al., 2020; Rizzuto et al., 2021; van Beest et al., 2023). For example, previous work found clear signals of white birch (*Betula papyrifera*) quality on moose habitat selection (Balluffi-Fry et al., 2020) from StDMs with predictive capacities (31%) comparable to those observed here. Given the diversity of species reported in caribou diet (Webber et al., 2022) and the absence of Fogo-specific diet composition data, we sampled more broadly, collecting multiple forage species from each site (mean = 3 species per site, range = 1 – 5). Still, we detected clear responses in selection for %N of *Cladonia* lichen in winter, and of vascular plant %N in summer. We expect that more accurate characterization of seasonal diets, more sampling sites across a wider range of environments, and clearer matches between animal space use and plant samples would explain even more variation in our models.

Forage quality is influenced by spatial heterogeneity in elemental distributions, which are in turn structured by abiotic characteristics such as underlying geology (Augusto et al., 2017; Morford et al., 2011), topography (Weintraub et al., 2015), and hydrology (Houlton & Morford, 2015). Forage quality differed both widely across the landscape scale and at the local scale. While our analyses suggest the Fogo Island landscape is overall nitrogen-poor, even small differences in forage quality can be ecologically consequential. We found fine-scale responses of caribou to relative forage quality at the individual step level, indicating that they may be able to detect and select for even slight variation in elemental content.

Caribou selecting areas with higher nitrogen at the step scale reflects individual-level behavioural decisions at a two-hour interval. When scaled up, however, this individual behaviour could have landscape-level consequences for elemental distribution. Animal-derived inputs are known to create hotspots of elements across landscapes (Bump et al., 2009; Ferraro et al., 2022; Monk, 2024), and in northern ecosytems these hotspots can increase forage quality over long periods (Barthelemy et al., 2024; Danell et al., 2002). Ungulates are known to locate and repeatedly use areas previously fertilized by urine (Day & Detling, 1990; Steinauer & Collins, 2001), fomenting a feedback in which animal aggregation and excretion may help generate and reinforce the very forage-quality patterns that animals subsequently select. Fine-scale individual behaviour could therefore go on to create landscape-level associations between population and elemental distributions.

Despite being low in nitrogen overall, forage on the caribou winter range appeared to be slightly higher in nitrogen relative to the broader landscape. Caribou are not necessarily the sole or primary cause of this nitrogen enrichment, as caribou may simply select existing higher-quality locations in which to spend their time overwinter. The spatial pattern does however differentiate two questions: how do relatively enriched areas *form* in a nitrogen-limited system characterized by low atmospheric deposition (Cheng et al., 2022) and minimal soil development, but also how are said areas *maintained* through time? One possible mechanism is animal-mediated transport of nitrogen across the landscape. The preliminary evidence we present here, when other caribou research, suggests that caribou movement and deposition contribute to spatially structured nitrogen patterns. Woodland caribou are more aggregated in winter than summer (Peignier et al., 2019), and summer and winter intensity of use at a given location were negatively correlated. We estimate that Fogo Island caribou release 2.2 kg/km^2^ of nitrogen via winter catabolism. This input is approximately 11% of atmospheric nitrogen deposition, a non-trivial input in a system where baseline nitrogen availability is low. In such systems, even modest animal-derived nitrogen inputs can have outsized ecosystem consequences, reinforcing localized nutrient hotspots (Barthelemy et al., 2018; Ferraro et al., 2024). Fogo caribou might have initially selected winter locations that were already higher in nitrogen, but by virtue of aggregating and catabolizing muscle in these areas, they could maintain if not magnify this spatial pattern.

From the animal perspective, concentrated nitrogen inputs could constitute an evolutionary strategy to mitigate seasonal nutrient deficits, rather than simply an accidental byproduct of spatial behaviour. Nitrogen accumulated during summer is later mobilized through winter catabolism (Barboza & Parker, 2008). At the population level, the nitrogen stored in muscle tissue is amassed from a wider spatial area when caribou are dispersed and nitrogen is most available. Because woodland caribou then aggregate on winter grounds, nitrogen releases are concentrated rather than diffuse. Repeated aggregation has been shown in other systems to generate persistent elemental hotspots, including grazing lawns, which can alter nutrient availability and forage quality at the landscape scale (Augustine et al., 2003). In nitrogen-poor landscapes, this spatial concentration of nitrogen may help reinforce localized nutrient availability while potentially buffering winter nutritional constraints. Further, if lichen takes up nitrogen deposited in urine, this improved forage quality could strengthen the benefits of site fidelity to winter grounds and reinforce the cycle. In this way, winter aggregation and catabolic nitrogen release may function as a coupled mechanism that both shapes the nutrient landscape experienced during subsequent foraging, and mitigates winter nitrogen scarcity (Box 1).

Importantly, the cyclicity between landscape distributions and the decisions animals make on the landscape makes it difficult to identify discrete cause and effect. This cyclicity is an inherent part of the system and, in our view, a feature, not a bug. Do caribou (a) select sites in winter because the lichen there is high in nitrogen, or (b) is the lichen high in nitrogen there because it is a high use caribou area? We are unable to distinguish between these given our data, but the feedback mechanism we propose would suggest both (a) and (b) are simultaneously plausible. There are additional steps that could be undertaken to demonstrate more mechanistic direct evidence of nutrient transport (Box 1), but for the purposes of our question, the emergent patterns in this system align with our expectations.

### Conclusion

Winter is the limiting season for most wildlife across the Boreal forest. We suggest that the resilience of caribou to winter severity might in part be attributed to the use of two different methods in concert: the movement to areas of relatively nitrogen-rich food and the presence of the enhanced forage quality via nitrogen deposition from catabolized material. Nitrogen deposition in urine-enriched winter forage could be understood as a form of caching through an external “root cellar,” where caribou transfer bodily stores of nitrogen into lichen and soil to be reused in future years. The extra-corporeal caching process we propose would be strongest with consistent space use and site fidelity across seasons, but there are of course other drivers of herbivore space use; predation risk is often suggested to restrict caribou more than forage availability (Briand et al., 2009). We do not suggest that the deposition of catabolized nitrogen is the origin or primary driver of winter site fidelity or sociality, but rather that it may be both the result of these behaviours and contribute to the maintenance of these behaviours. Catabolism-mediated nutrient transport via seasonal changes in animal space use and physiology could be an underappreciated process in other ecosystems, influencing both distribution of elements but also feeding back into the behaviour of the animal vectors themselves.

#### Box 1

##### Conceptual basis and future work required to demonstrate adaptive nitrogen transport and accumulation by woodland caribou.

The interaction between nutrient transport and forage selection we describe here could constitute a form of zoogeochemical niche construction (ZNC; Ferraro et al., 2025). ZNC is a complex phenomenon requiring several processes. Niche construction theory in the broad sense involves organisms modifying the selective environment for themselves or other species (Laland et al., 2016; Odling-Smee et al., 1996). Two criteria define ZNC specifically: (1) non-random geochemical modification of the environment by the focal species, and (2) modified selective pressure on the recipient organism caused by this chemical modification. Once identified, (3) an evolutionary response in the recipient organism can then be assessed (Ferraro et al., 2025). Here, we explore the potential for aggregated nitrogen deposition on winter grounds—arising from caribou catabolism during nutritionally restrictive periods—to represent an instance of ZNC. Our study provides evidence supporting the first criterion and outlines a plausible mechanism for the second, though testing for evolutionary responses lies beyond the scope of this study. Below, we identify the additional research questions necessary to more definitively establish these causal relationships.

First, we base our transport mechanism on the seasonal shift from nitrogen surplus to deficit that occurs in caribou metabolism from summer to winter. This physiological process has been documented across populations (Barboza et al., 2018; Barboza & Parker, 2008; Parker et al., 2005), though it might not be as strong for some herds than for island populations in which forage is more limiting (Adamczewski et al., 1987). While earlier work tended to involve lethal procedures to quantify bodily stores of nitrogen (Huot, 1989), less invasive processes including stable isotope analyses have been developed (Gustine et al., 2012) to document nitrogen flux within caribou throughout the year. Fogo Island appears to be nitrogen-limited, so whether Fogo caribou experience the same range of forage quality as other populations remains to be seen. Our first open question is thus *(1) What does the annual nitrogen budget of caribou within a specific system look like, including the magnitude of muscle tissue accretion in summer and lean mass loss over winter?*

Isotopic analyses could also help with a second question: *(2) Is the nitrogen deposited by caribou during winter taken up by forage and made nutritionally available to caribou once more?* Animal-derived inputs, including urine, should have a signature of d15N enrichment (Gustine et al., 2012) that could be used to identify animal-sourced N in plants and lichen. Experimental simulations of parturition on Fogo did find that urea deposited at point sites was taken up by several species, particularly so by *Cladonia* lichens (Ferraro et al., 2024), but this experiment took place in summer. The capacity for vascular and non-vascular plants to assimilate nutrients deposited on snow at sub-zero temperatures is less clear (but see Preston et al., 1990). Lichens are able to rapidly become metabolically active during winter, unlike vascular plants that are limited by seasonal dormancy (Bjerke, 2010; Lange, 2003) and may be better able to take up nutrients through winter (Kappen, 1993), but the rate of this intake and latency until it is metabolised into nutritive compounds for caribou to consume is unknown.

(3) *Do caribou preferentially use sites enriched with urine-derived nitrogen?* In grassland systems, bison repeatedly select previously urinated patches, likely tracking localized increases in forage quality (Day & Detling, 1990; Steinauer & Collins, 2001). Whether caribou exhibit similar behavior remains unknown, as well as the latency before which they return. If caribou are able to detect and repeatedly forage within urine enriched patches, such behavior would strengthen the case for a feedback loop between element deposition and habitat use.

Finally, the nitrogen ZNC we propose is the result of a behavioural phenotype involving consistent seasonal shifts in diet, space use, sociality, and site fidelity. As such, a key unanswered question is *(4) Is this socio-spatial behavioural phenotype adaptive, or do other selective pressures discourage this behaviour?* The landscape distributions of caribou and nitrogen broadly aligned with the mechanism we propose. However, if nitrogen deposition does improve winter forage, the positive feedback to facilitate this behaviour’s persistence would require individual-level fitness benefits. Improved body condition in spring, being the proximal mechanism, could be one fitness proxy, while improved calf survival or probability of parturition would be a more direct reproductive success metric.

## Supporting information

SupportingInformationA-D

SupportingInformationE

## Acknowledgements

Data collection, analysis, and writing for this study took place on the ancestral homelands of the Mi’kmaq and Beothuk. Thank you to Christina Prokopenko, Julie Turner, and Alec Robitaille for their invaluable guidance and assistance with models as part of the iSSA guild, and to Sam Goguen for assistance in the field. We would also like to thank Gail and Ferg Penton and Fraser Carpenter for their kindness and generosity in supporting us during our time on Fogo Island.

## Author contributions

JGH, KMF, and EVW conceived the ideas and designed methodology; JGH, KMF, and DA collected the data; JGH, KMF, and JMK analysed the data; JGH led the writing of the manuscript; KMF, AEL, JMK, SJL, and QMRW edited and revised the manuscript extensively. All authors contributed critically to the drafts and gave final approval for publication.

## Statement on Inclusion

This research took place in the same province as the primary author’s institution within Canada. We worked closely with local residents of Fogo Island in the field, and provided opportunities for co-authorship to undergraduate and early-career collaborators.

## Data availability statement

Data and code for StDM analyses and iSSA are available at https://doi.org/10.5281/zenodo.18760996 (Hendrix et al., 2026a) and https://doi.org/10.5281/zenodo.18763026 (Hendrix et al., 2026b), respectively.

