## SupportingInformationA-D for "Shifting forage selection subsidizes seasonal resource scarcity"

### Appendix A: Stoichiometric distribution modelling details

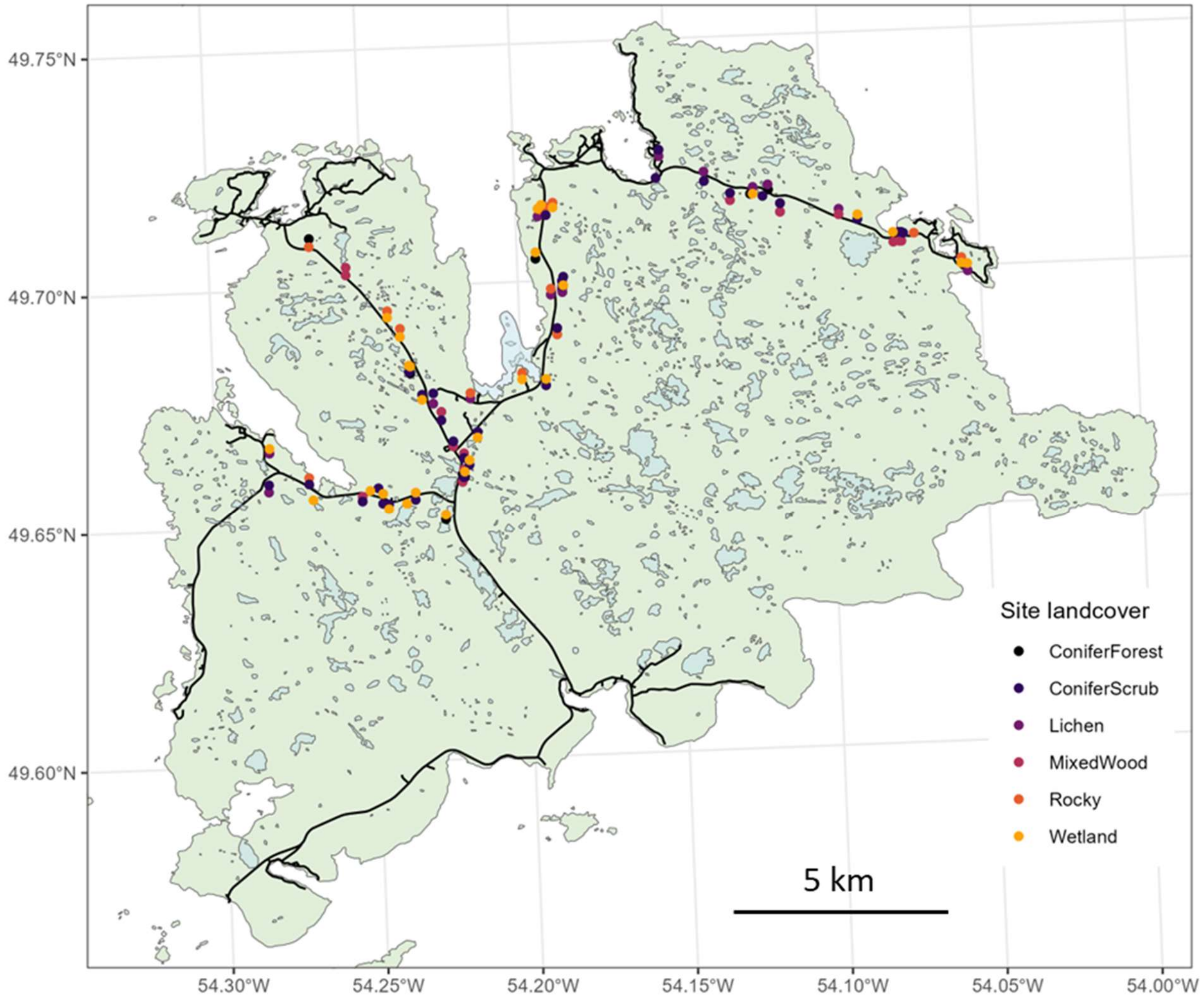


Figure A1. Forage sampling locations on Fogo Island, NL, between June 11 – 14 2022. Sites were located within 500m but more than 250m from the road for access. Colour of point represents the land cover classification from the provincial land classification system (Integrated Informatics Inc., 2014).

**Stoichiometric distribution models for Cladonia and average vascular %N**

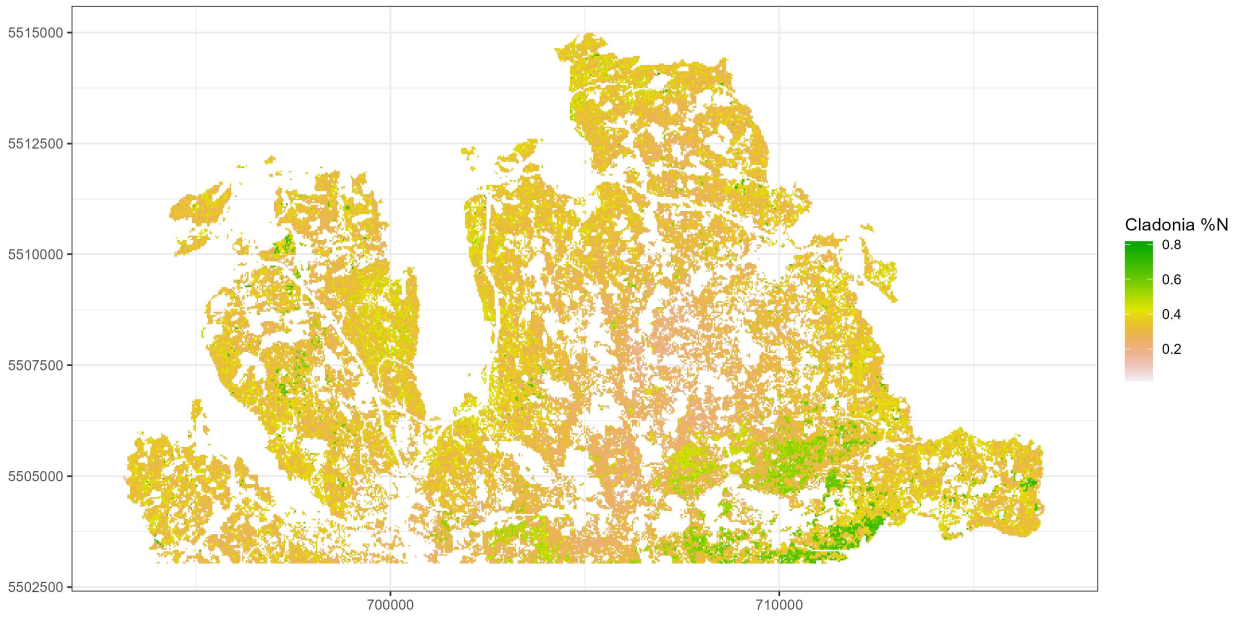


Figure A2. Predicted %N content in *Cladonia* lichen across northern half of Fogo Island, NL. Gaps in data reflect land cover categories (e.g. water, roads) where forage samples were not collected.

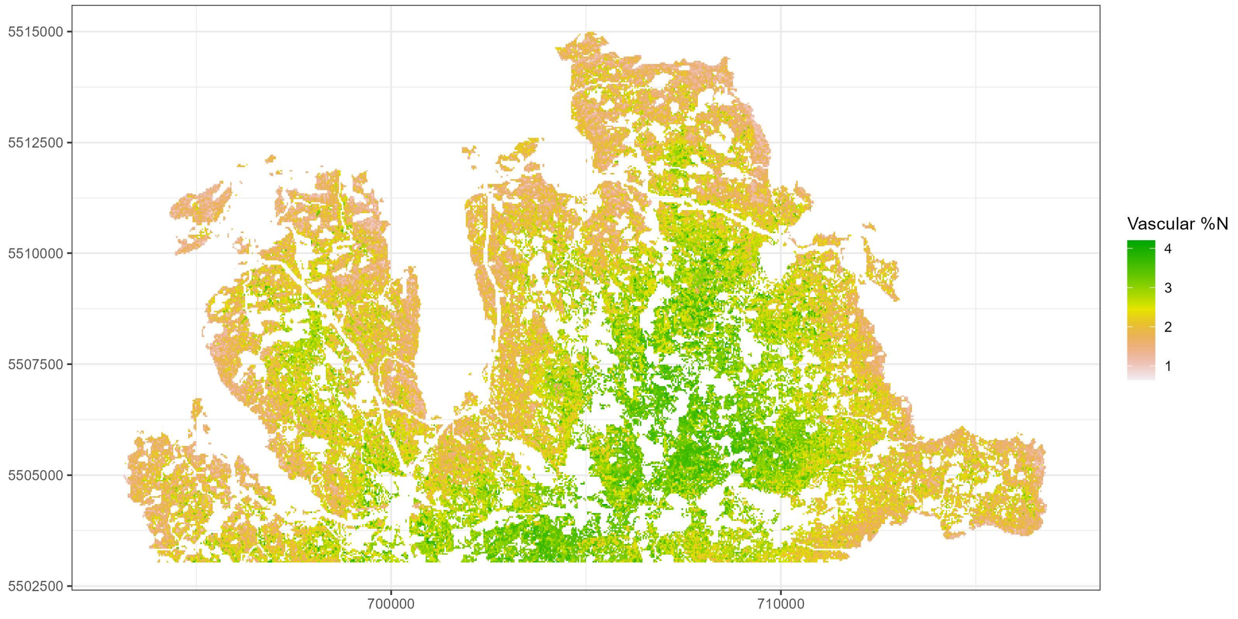


Figure A3. Average predicted %N content of seven vascular plant species across northern half of Fogo Island, NL. Gaps in data reflect land cover categories (e.g. water, roads) where forage samples were not collected.

#### Predictive models of % nitrogen content for each species

Table A1. Model summaries for beta regression predictive models of % nitrogen content for forage species on Fogo Island, NL. "CFS" refers to Canadian Forest Service land cover classes as a categorical predictor derived from Hermosilla et al (2022). "NL" reflects provincial land cover classification by the Newfoundland and Labrador SDSS (Integrated Informatics, 2014).
* The model for dwarf birch was abridged so that convergence was achieved, with CFS land cover and terrain ruggedness removed as predictors.

| Alder | $R^2$ = 63.5%, n samples = 111, n sites = 27 | | |
|---|---|---|---|
| | Estimate ± SE | *z* | *p* |
| Intercept | -5.979 ± 0.725 | -8.25 | < 0.001 |
| CFS Mixedwood | 0.0586 ± 0.041 | 1.44 | 0.15 |
| CFS Shrubs | 0.183 ± 0.086 | 2.12 | 0.03 |
| CFS Wetland | -0.0497 ± 0.064 | -0.77 | 0.44 |
| NL ConiferScrub | 0.4701 ± 0.079 | 5.94 | < 0.001 |
| NL Lichen | 0.4491 ± 0.126 | 3.56 | < 0.001 |
| NL MixedWood | 0.4085 ± 0.085 | 4.80 | < 0.001 |
| NL Rocky | 0.5981 ± 0.128 | 4.66 | < 0.001 |
| NL Wetland | 0.3344 ± 0.08 | 4.16 | < 0.001 |
| Distance to coast (km) | 0.3528 ± 0.055 | 6.44 | < 0.001 |
| NDVI | 0.0001 ± 0.00004 | 3.23 | 0.00 |
| Elevation (m) | -0.0031 ± 0.001 | -3.53 | < 0.001 |
| Slope | -0.0519 ± 0.014 | -3.83 | < 0.001 |
| cos(Aspect) | 0.0314 ± 0.029 | 1.08 | 0.28 |
| Topographic position index | -0.7529 ± 0.134 | -5.62 | < 0.001 |
| Terrain ruggedness index | 0.1681 ± 0.055 | 3.06 | 0.00 |

| Blueberry | $R^2$ = 44.1%, n samples = 147, n sites = 37 | | |
|---|---|---|---|
| | Estimate ± SE | *z* | *p* |
| Intercept | -2.838 ± 0.761 | -3.73 | < 0.001 |
| CFS Coniferous | 0.2184 ± 0.165 | 1.32 | 0.19 |
| CFS Mixedwood | 0.0504 ± 0.212 | 0.24 | 0.81 |
| CFS Shrubs | 0.4006 ± 0.157 | 2.56 | 0.01 |
| CFS Wetland | 0.0895 ± 0.181 | 0.49 | 0.62 |
| NL Lichen | -0.3433 ± 0.068 | -5.05 | < 0.001 |
| NL MixedWood | 0.0276 ± 0.08 | 0.34 | 0.73 |
| NL Rocky | -0.4099 ± 0.107 | -3.83 | < 0.001 |
| NL Wetland | -0.241 ± 0.139 | -1.73 | 0.08 |
| Distance to coast (km) | 0.0212 ± 0.083 | 0.26 | 0.80 |
| NDVI | 0.00004 ± 0.00005 | -0.93 | 0.35 |
| Elevation (m) | 0.0011 ± 0.002 | 0.63 | 0.53 |
| Slope | -0.0174 ± 0.018 | -0.94 | 0.35 |
| cos(Aspect) | 0.0592 ± 0.048 | 1.23 | 0.22 |
| Topographic position index | 0.0217 ± 0.099 | 0.22 | 0.83 |

| | | | |
|---|---|---|---|
| Terrain ruggedness index | -0.0215 ± 0.071 | -0.30 | 0.76 |

| *Cladonia* | $R^2$ = 23.9%, n samples = 248, n sites = 63 | | |
|---|---|---|---|
| | Estimate ± SE | *z* | *p* |
| Intercept | -4.616 ± 0.6478 | -7.125 | < 0.001 |
| CFS Coniferous | -0.09801 ± 0.09143 | -1.072 | 0.28 |
| CFS Shrubs | -0.095 ± 0.07331 | -1.296 | 0.2 |
| CFS Wetland | -0.1709 ± 0.07163 | -2.386 | 0.017 |
| NL ConiferForest | 0.5404 ± 0.2227 | 2.426 | 0.015 |
| NL ConiferScrub | -0.01571 ± 0.1795 | -0.088 | 0.93 |
| NL Lichen | -0.2276 ± 0.1847 | -1.232 | 0.22 |
| NL MixedWood | -0.3452 ± 0.2121 | -1.627 | 0.1 |
| NL Rocky | -0.2199 ± 0.1949 | -1.128 | 0.26 |
| NL Wetland | 0.04212 ± 0.1785 | 0.236 | 0.81 |
| Distance to coast (km) | -0.1201 ± 0.0787 | -1.525 | 0.13 |
| NDVI (m) | -0.04695 ± 0.03491 | -1.345 | 0.18 |
| Elevation (m) | 0.001345 ± 0.001793 | 0.75 | 0.45 |
| Slope | 0.0227 ± 0.01045 | 2.172 | 0.03 |
| cos(Aspect) | 0.04727 ± 0.0422 | 1.12 | 0.26 |
| Topographic position index | -0.008529 ± 0.1338 | -0.064 | 0.95 |
| Terrain ruggedness index | -0.13 ± 0.04991 | -2.604 | 0.009 |

| Crowberry | $R^2$ = 19.9%, n samples = 190, n sites = 48 | | |
|---|---|---|---|
| | Estimate ± SE | *z* | *p* |
| Intercept | -3.816 ± 0.595 | -6.42 | < 0.001 |
| CFS Coniferous | -0.19 ± 0.098 | -1.95 | 0.05 |
| CFS Shrubs | -0.0052 ± 0.082 | -0.06 | 0.95 |
| CFS Wetland | -0.2288 ± 0.112 | -2.04 | 0.04 |
| NL ConiferScrub | -0.5765 ± 0.264 | -2.19 | 0.03 |
| NL Lichen | -0.0173 ± 0.161 | -0.11 | 0.91 |
| NL MixedWood | -0.2613 ± 0.225 | -1.16 | 0.25 |
| NL Rocky | -0.0887 ± 0.214 | -0.41 | 0.68 |
| NL Wetland | 0.0772 ± 0.168 | 0.46 | 0.65 |
| Distance to coast (km) | -0.0253 ± 0.135 | -0.19 | 0.85 |
| NDVI | -0.00006 ± 0.00003 | -1.71 | 0.09 |
| Elevation (m) | 0.0023 ± 0.004 | 0.61 | 0.54 |
| Slope | -0.0365 ± 0.019 | -1.92 | 0.06 |
| cos(Aspect) | 0.023 ± 0.061 | 0.38 | 0.70 |
| Topographic position index | 0.1274 ± 0.208 | 0.61 | 0.54 |
| Terrain ruggedness index | 0.1557 ± 0.07 | 2.23 | 0.03 |

| Deergrass | $R^2$ = 0.364, n samples = 76, n sites = 19 | | |
|---|---|---|---|
| | Estimate ± SE | *z* | *p* |
| Intercept | -0.243 ± 1.186 | -0.21 | 0.84 |

| | | | |
|---|---|---|---|
| CFS Coniferous | 0.2375 ± 0.157 | 1.52 | 0.13 |
| CFS Shrubs | 0.1569 ± 0.149 | 1.06 | 0.29 |
| CFS Wetland | 0.0973 ± 0.12 | 0.81 | 0.42 |
| NL Rocky | 0.699 ± 0.302 | 2.32 | 0.02 |
| NL Wetland | 0.1991 ± 0.107 | 1.85 | 0.06 |
| Distance to coast (km) | 0.3058 ± 0.106 | 2.90 | 0.004 |
| NDVI | -0.0002 ± 0.00008 | -3.07 | 0.002 |
| Elevation (m) | -0.0094 ± 0.003 | -2.70 | 0.01 |
| Slope | -0.0094 ± 0.016 | -0.61 | 0.54 |
| cos(Aspect) | 0.0797 ± 0.049 | 1.61 | 0.11 |
| Topographic position index | 0.2308 ± 0.15 | 1.54 | 0.12 |
| Terrain ruggedness index | 0.0248 ± 0.059 | 0.42 | 0.68 |

| Dwarf birch* | $R^2$ = 37.9%, n samples = 60, n sites = 15 | | |
|---|---|---|---|
| | Estimate ± SE | *z* | *p* |
| Intercept | -5.818 ± 1.072 | -5.43 | < 0.001 |
| NL Wetland | 0.2231 ± 0.115 | 1.95 | 0.05 |
| Distance to coast (km) | 0.2871 ± 0.125 | 2.30 | 0.02 |
| NDVI | 0.0001 ± 0.00006 | 2.13 | 0.03 |
| Elevation (m) | 0.0087 ± 0.002 | 3.67 | < 0.001 |
| Slope | -0.083 ± 0.018 | -4.73 | < 0.001 |
| cos(Aspect) | 0.6114 ± 0.139 | 4.41 | < 0.001 |
| Topographic position index | 0.1856 ± 0.086 | 2.17 | 0.03 |

| Kalmia | $R^2$ = 31.6%, n samples = 334, n sites = 85 | | |
|---|---|---|---|
| | Estimate ± SE | *z* | *p* |
| Intercept | -3.274 ± 0.346 | -9.46 | < 0.001 |
| CFS Coniferous | 0.0122 ± 0.054 | 0.22 | 0.82 |
| CFS Mixedwood | 0.0911 ± 0.072 | 1.27 | 0.21 |
| CFS Shrubs | 0.0098 ± 0.049 | 0.20 | 0.84 |
| CFS Wetland | -0.0398 ± 0.044 | -0.91 | 0.36 |
| NL ConiferForest | -0.04 ± 0.135 | -0.30 | 0.77 |
| NL ConiferScrub | -0.0853 ± 0.075 | -1.13 | 0.26 |
| NL Lichen | -0.2283 ± 0.075 | -3.03 | 0.002 |
| NL MixedWood | -0.0202 ± 0.076 | -0.27 | 0.79 |
| NL Rocky | -0.2158 ± 0.09 | -2.41 | 0.02 |
| NL Wetland | -0.1273 ± 0.084 | -1.52 | 0.13 |
| Distance to coast (km) | -0.0001 ± 0 | -1.71 | 0.09 |
| NDVI | 0 ± 0 | -2.28 | 0.02 |
| Elevation (m) | 0.0004 ± 0.001 | 0.51 | 0.61 |
| Slope | 0.0114 ± 0.006 | 1.89 | 0.06 |
| cos(Aspect) | 0.0457 ± 0.017 | 2.73 | 0.01 |
| Topographic position index | 0.1522 ± 0.055 | 2.75 | 0.01 |
| Terrain ruggedness index | -0.1241 ± 0.026 | -4.84 | < 0.001 |

| Lingonberry | $R^2$ = 47.1%, n samples = 177, n sites = 44 | | |
|---|---|---|---|
| | Estimate ± SE | *z* | *p* |
| Intercept | -4.268 ± 0.836 | -5.10 | < 0.001 |
| CFS Coniferous | -0.0037 ± 0.09 | -0.04 | 0.97 |
| CFS Mixedwood | 0.5436 ± 0.135 | 4.04 | < 0.001 |
| CFS Shrubs | -0.0597 ± 0.078 | -0.76 | 0.45 |
| CFS Wetland | -0.0254 ± 0.074 | -0.35 | 0.73 |
| NL ConiferScrub | 0.0693 ± 0.093 | 0.75 | 0.45 |
| NL Lichen | -0.1954 ± 0.09 | -2.17 | 0.03 |
| NL MixedWood | -0.0046 ± 0.086 | -0.05 | 0.96 |
| NL Rocky | -0.2468 ± 0.119 | -2.08 | 0.04 |
| Distance to coast (km) | 0.2147 ± 0.052 | 4.15 | < 0.001 |
| NDVI | .00002 ± 0.0005 | -0.46 | 0.65 |
| Elevation (m) | -0.0058 ± 0.001 | -4.54 | < 0.001 |
| Slope | 0.0351 ± 0.008 | 4.28 | < 0.001 |
| cos(Aspect) | 0.0385 ± 0.026 | 1.47 | 0.14 |
| Topographic position index | 0.4172 ± 0.089 | 4.67 | < 0.001 |
| Terrain ruggedness index | -0.2385 ± 0.04 | -6.00 | < 0.001 |

Integrated Informatics Inc. (2014). *Sustainable Development & Strategic Science Branch land cover classification.* Sustainable Development and Strategic Science, Government of Newfoundland and Labrador.

Hermosilla, T., Wulder, M. A., White, J. C., & Coops, N. C. (2022). Land cover classification in an era of big and open data: Optimizing localized implementation and training data selection to improve mapping outcomes. *Remote Sensing of Environment*, *268*, 112780. https://doi.org/10.1016/j.rse.2021.112780

**Correlation matrix of species % nitrogen content**

Table A2. Pairwise correlations in average % nitrogen of forage samples from n = 108 sites. Note that some species did not occur in sympatry, or were only observed once or twice at the same site; in such cases, a correlation cannot be drawn between such species (or is exactly -1, i.e. Cladonia vs. dwarf birch).

| % N **from sample sites (n = # of co-occurrences)** | crow berry | sheep laurel | lingon berry | alder | blue berry | deer grass | dwarf birch |
|---|---|---|---|---|---|---|---|
| Cladonia | 44% n = 45 | 44% n = 57 | 36% n = 34 | NA n = 0 | 41% n = 45 | NA n = 1 | -100% n = 2 |
| crowberry | | 18% n = 43 | 25% n = 30 | NA n = 0 | 38% n = 18 | NA n = 0 | NA n = 1 |
| sheep laurel | | | 47% n = 38 | 12% n = 8 | 57% n = 27 | 49% n = 7 | -26% n = 7 |
| lingonberry | | | | 37% n = 5 | 44% n = 18 | NA n = 0 | NA n = 0 |
| alder | | | | | -42% n = 8 | NA n = 0 | -51% n = 4 |
| blueberry | | | | | | NA n = 1 | NA n = 1 |
| deergrass | | | | | | | -33% n = 12 |

Table A3. Correlations between predicted %N content of forage species in stoichiometric distribution models on Fogo Island, NL. Note that some species were more likely to co-occur in space than others, so the number of observations leading to each pairwise correlation estimate varied.

| % N correlations **From StDMs** | crow berry | sheep laurel | lingon berry | alder | blue berry | deer grass | dwarf birch |
|---|---|---|---|---|---|---|---|
| Cladonia | 8% | 49% | -9% | -48% | -1% | -31% | -25% |
| crowberry | | -19% | -58% | -29% | -54% | 43% | 10% |
| sheep laurel | | | 44% | -29% | 48% | -12% | 12% |
| lingonberry | | | | 36% | 44% | 10% | 40% |
| alder | | | | | 43% | 8% | 49% |
| blueberry | | | | | | -26% | 41% |
| deergrass | | | | | | | 34% |

Table A4. Average percent carbon content and C:N ratio of forage species samples collected on Fogo Island, NL, June 11 –14 2022, along with model pseudo-$R^2$ for predictive models. C:N ratio was inverted to N:C ratio prior to modelling so that the same beta regression model as for %C and %N could be applied to the ratio between 0 and 1; when predicting ratio across the rest of the study area, this was re-converted back to C:N for visualization purposes. The same model was applied to each species* and included Canadian Forest Service land cover, provincial land cover, NDVI, elevation (m), slope (%), aspect (°), distance to coast (m), topographic position index, and terrain ruggedness index.
* CFS land cover and terrain ruggedness were removed from the dwarf birch models for convergence.

| Species | *n* samples | % carbon ± SE | Pseudo-$R^2$ of carbon model (%) | C:N ratio ± SE | Pseudo-$R^2$ of C:N model (%) |
|---|---|---|---|---|---|
| Alder | 111 | 49.47 ± 0.11 | 53.5 | 12.92 ± 0.24 | 63.8 |
| Blueberry | 147 | 50.68 ± 0.17 | 16.1 | 21.44 ± 0.66 | 38.4 |
| *Cladonia* | 247 | 44.49 ± 0.26 | 12.9 | 152.27 ± 2.82 | 28.3 |
| Crowberry | 189 | 55.84 ± 0.32 | 8.7 | 77.16 ± 1.29 | 21.2 |
| Deergrass | 76 | 45.75 ± 0.11 | 29.4 | 24.44 ± 0.79 | 25.1 |
| Dwarf birch | 60 | 49.43 ± 0.19 | 33.8 | 16.92 ± 0.49 | 33.8 |
| Lingonberry | 176 | 51.88 ± 0.18 | 39 | 81.95 ± 1.45 | 42.7 |
| Sheep laurel | 332 | 53.87 ± 0.23 | 11.4 | 43.58 ± 0.5 | 31.4 |

Species-specific parameter estimates for models predicting %C and C:N ratio, analogous to Table A1, can be obtained in code provided at https://doi.org/10.5281/zenodo.18760996

#### Appendix B: Winter subsamping at select sites

Our StDM surfaces are based on data collected in June. We did revisit ten sites in winter for comparison, though ten sites were insufficient to generate an StDM. As we were unable to relocate the exact same 50 x 50cm quadrat from our summer sites, we do not compare specific locations across seasons, but rather took the average of each species samples within season (n = 4 subsamples x 10 sites = 40 per species) to look at general trends. The seasonal comparisons demonstrate that *Cladonia* nitrogen content is relatively consistent throughout the year as expected (Chapin et al., 1980). Converse to our expectations our vascular plant species samples did not decline as much as might be expected (Fig B1). Most of the vascular plants sampled retain their foliage through the winter. Only the deciduous species (blueberry) illustrated clear seasonal differences, as the winter blueberry samples consisted of twigs and buds, not leaves as in summer.

Nitrogen concentration in broadleaf evergreen leaves, e.g., Labrador tea, tends to decrease through the growing season into winter in their first year of emergence. Leaves on plants for their second or third year tend to remain at that same low level of nitrogen in subsequent growing seasons and change minimally throughout the annual cycle until they senesce (Prudhomme, 1983). Phenolics and tannin concentrations tend to parallel this change, with the highest production of anti-herbivore secondary metabolites when leaves are richest in N and most attractive to herbivores, investing less in defense when leaves are not as appealing (Happe et al., 1990). While secondary metabolites may not explain the shift in caribou diet from summer to winter, caribou are known to focus on lichen overwinter and graminoids and shrubs in summer (Webber et al., 2022), and we showed here that caribou selected for higher %N in vascular in summer and %N in *Cladonia* in winter. We also note that average vascular % nitrogen and *Cladonia* % nitrogen were negatively correlated (r = - 0.058), though when looking at pairwise species comparisons, *Cladonia* was not as much an outlier as we had anticipated (Appendix A Table A2). Though this is a weak negative relationship, it does indicate a weak correlation between vascular plant and *Cladonia* quality, such that caribou are unlikely to simultaneously select for both. The best lichen and best vascular plants are either completely randomly associated with one another, or slightly non-overlapping, which may in part explain why seasonal ranges are similarly semi-distinct.

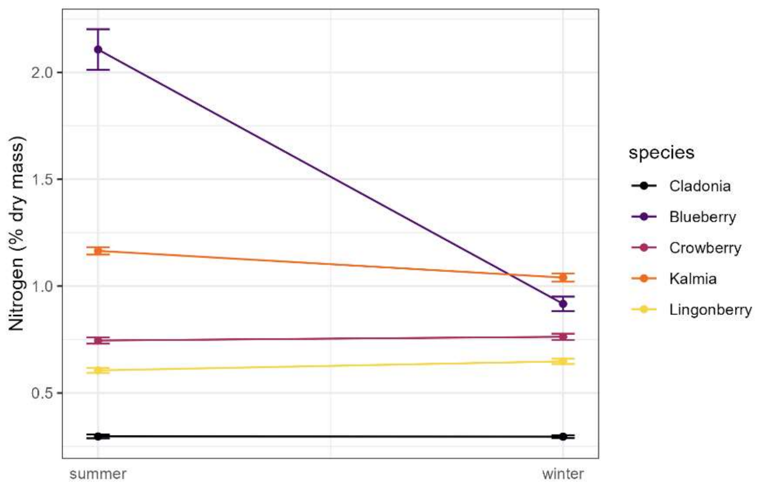


Figure B1. Change in % nitrogen content of five forage species on Fogo Island from summer to winter (n = 40 observations/species).

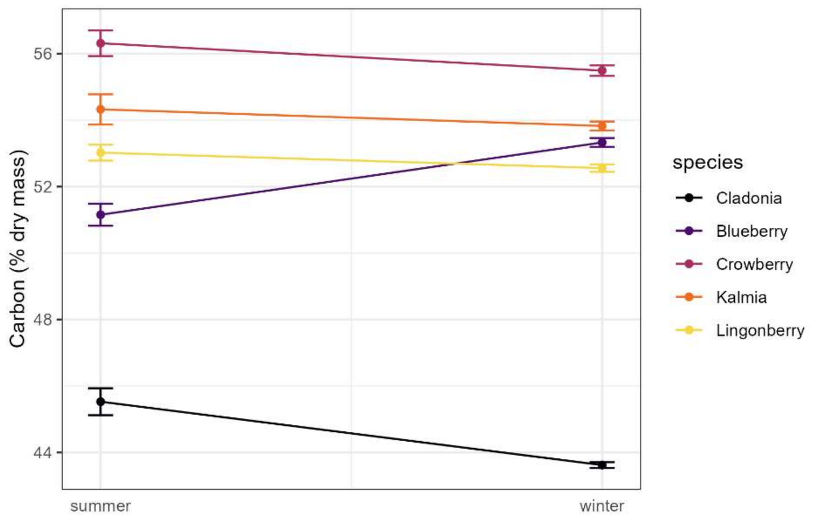


Figure B2. Change in % carbon content of five forage species on Fogo Island from summer to winter (n = 40 observations/species).

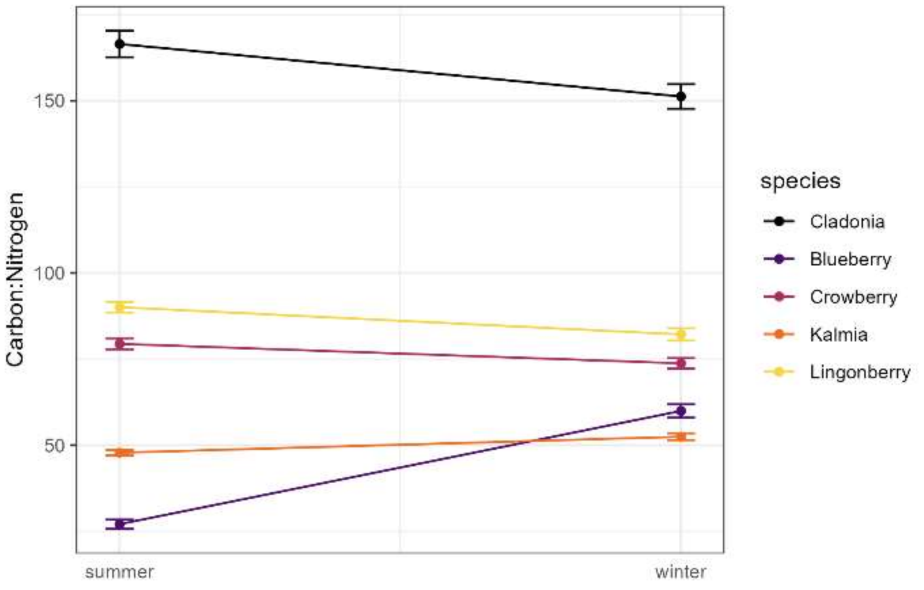


Figure B3. Change in carbon:nitrogen ratio in five forage species from summer to winter. Lower ratio values indicate higher quality forage. Contrary to expectations, not all vascular plant species declined in quality from summer to winter, and most were relatively consistent throughout the year.

Chapin, F. S., Johnson, D. A., & McKendrick, J. D. (1980). Seasonal movement of nutrients in plants of differing growth form in an Alaskan tundra ecosystem: Implications for herbivory. *Journal of Ecology*, *68*(1), 189–209. https://doi.org/10.2307/2259251

Happe, P. J., Jenkins, K. J., Starkey, E. E., & Sharrow, S. H. (1990). Nutritional quality and tannin astringency of browse in clear-cuts and old-growth forests. *The Journal of Wildlife Management*, *54*(4), 557–566. https://doi.org/10.2307/3809349

Prudhomme, T. I. (1983). Carbon allocation to antiherbivore compounds in a deciduous and an evergreen subarctic shrub species. *Oikos*, *40*(3), 344–356. https://doi.org/10.2307/3544307

Webber, Q. M. R., Ferraro, K. M., Hendrix, J. G., & Vander Wal, E. (2022). What do caribou eat? A review of the literature on caribou diet. *Canadian Journal of Zoology*, *100*(3), 197–207. https://doi.org/10.1139/cjz-2021-0162

### Appendix C: iSSA lineup validation protocol

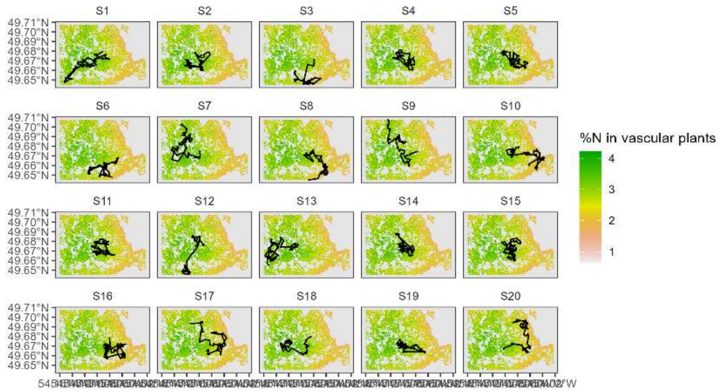


Figure S4. The lineup protocol presented to n = 9 observers, simulating summer movement data. One of these plots is a 5-day (n = 120 steps of 2h GPS data) segment of an individual caribou, while nineteen are simulations from the summer iSSA. [The real data segment is in panel S6].

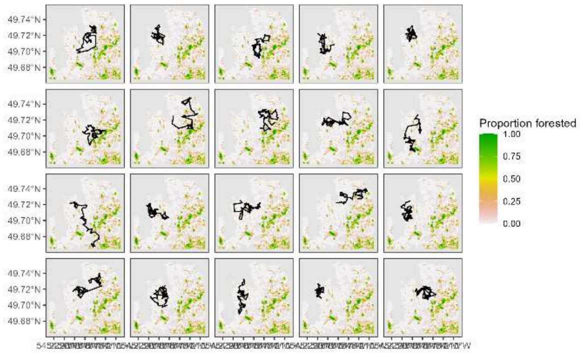


Figure S5. The lineup protocol presented to n = 9 observers, simulating winter movement data. One of these plots is a 5-day (n = 120 steps of 2h GPS data) segment of an individual caribou, while nineteen are simulations from the winter iSSA. [The real data segment is in panel W13].

**Appendix D: Relative selection strengths from iSSA**

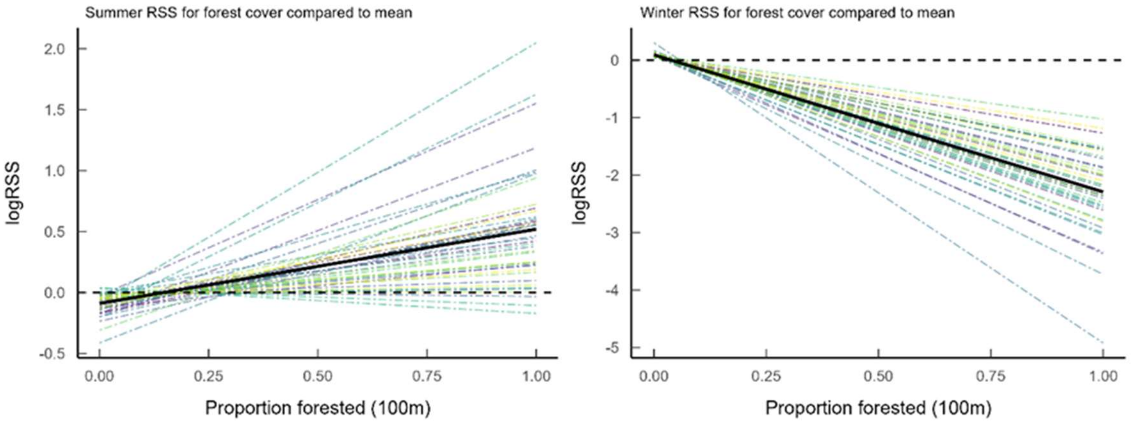


Figure D1. Relative selection strength for % forest cover in summer (June 16 – Aug 16) and winter (Jan 1 – Mar 1) by woodland caribou on Fogo Island, Newfoundland. Coloured dotted lines are each an individual caribou, with the solid black line representing the population-level response. RSS is calculated relative to the average % forest cover experienced by each individual (i.e. the x-intercept of each line).

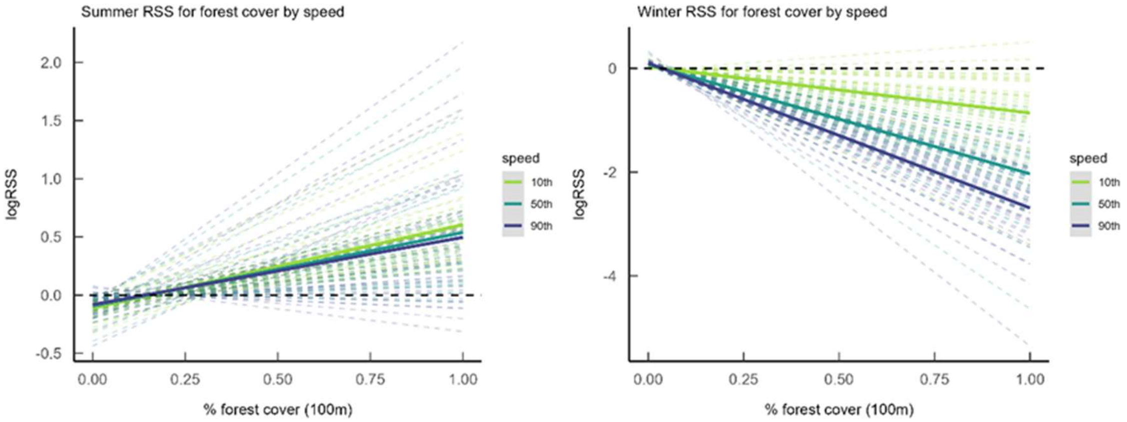


Figure D2. Relative selection strength for % forest cover as a function of movement rate in summer (June 16 – Aug 16) and winter (Jan 1 – Mar 1) by woodland caribou on Fogo Island, Newfoundland. Coloured dotted lines are each an individual caribou, with the bold solid lines representing the population-level response, for the $10^{th}$, $50^{th}$, and $90^{th}$ percentile of step lengths (= movement rate). These percentiles represent speeds of 4.6 or 4.1 m/h, 64 or 66 m/h, and 312 or 317 m/h in summer and winter, respectively. RSS is calculated relative to the average % forest cover experienced by each individual (i.e. the x-intercept of each line).

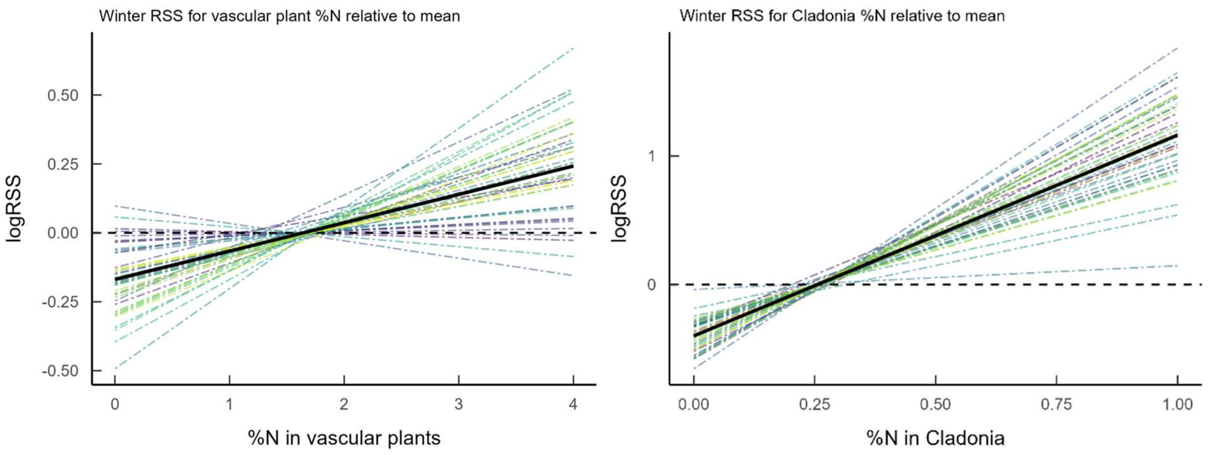


Figure D3. Individual RSS for two stoichiometric distribution models by woodland caribou on Fogo Island, NL. Winter refers to the period 1 January – 1 March. Relative selection strength represents the relative probability of selecting a given %N content relative to the average %N (where the line crosses the x-axis). Each coloured dotted line is an individual, with the solid black line showing the population average response.

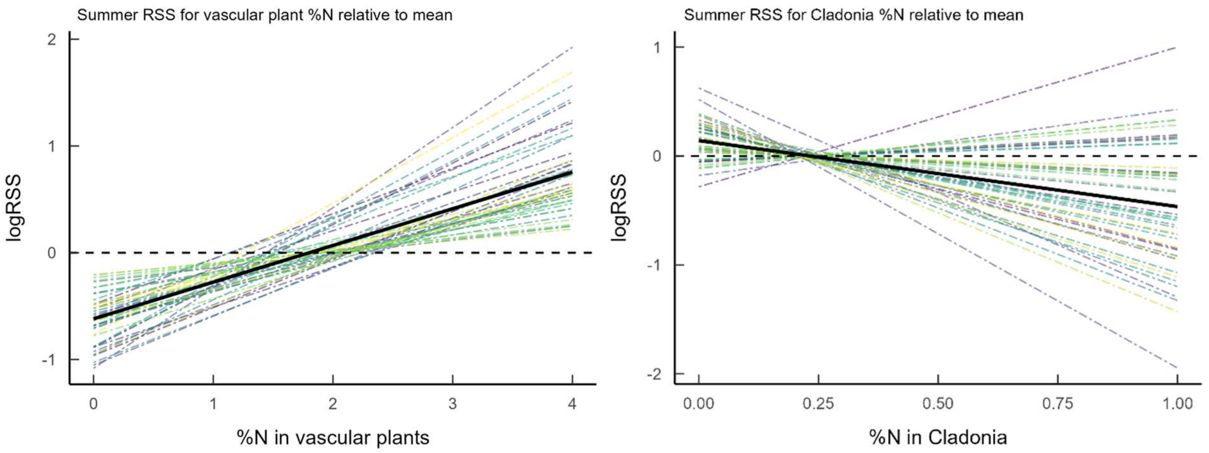


Figure D4. Individual RSS for two stoichiometric distribution models by woodland caribou on Fogo Island, NL. Summer refers to the period 16 June - 16 August. Relative selection strength represents the relative probability of selecting a given %N content relative to the average %N (where the line crosses the x-axis). Each coloured dotted line is an individual, with the solid black line showing the population average response.

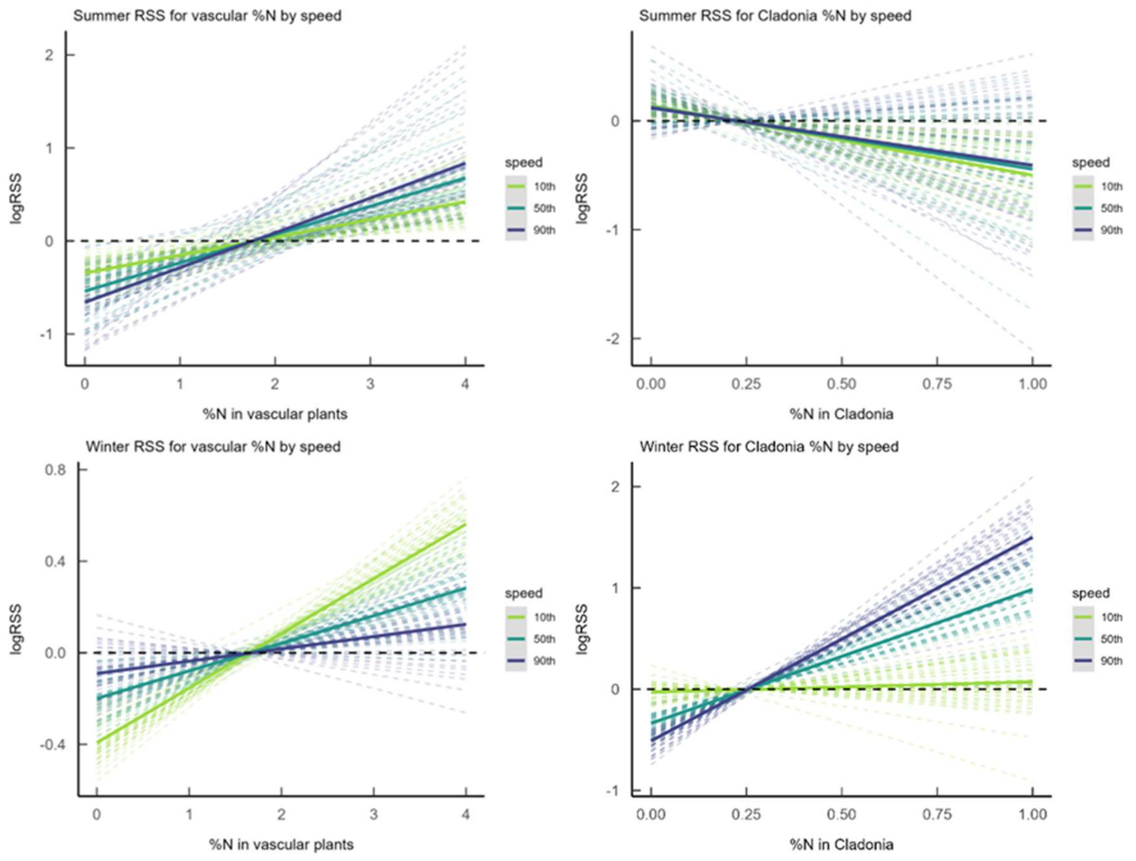


Figure D5. Relative selection strength for %N in forage as a function of movement rate in summer (June 16 – Aug 16) and winter (Jan 1 – Mar 1) by woodland caribou on Fogo Island, Newfoundland. Coloured dotted lines are each an individual caribou, with the bold solid lines representing the population-level response, for the 10$^{th}$, 50$^{th}$, and 90$^{th}$ percentile of step lengths (= movement rate). These percentiles represent speeds of 4.6 or 4.1 m/h, 64 or 66 m/h, and 312 or 317 m/h in summer and winter, respectively. RSS is calculated relative to the average %N experienced by each individual (i.e. the x-intercept of each line).
